# Frontoparietal dynamics and value accumulation in intertemporal choice

**DOI:** 10.1101/2020.08.05.237578

**Authors:** Qingfang Liu, Woojong Yi, Christian A. Rodriguez, Samuel M. McClure, Brandon M. Turner

**Author notes:** These authors contributed equally to this work (Qingfang Liu, Woojong Yi). This author is a senior author (Brandon M. Turner).

## Abstract

Intertemporal choice requires choosing between a smaller reward available after a shorter time delay and a larger reward available after a longer time delay. Previous studies suggest that intertemporal preferences are formed by generating a subjective value of the monetary rewards that depends on reward amount and the associated time delay. Neuroimaging results indicate that this subjective value is tracked by ventral medial prefrontal cortex (vmPFC) and ventral striatum. Subsequently, an accumulation process, subserved by a network including dorsal medial frontal cortex (dmFC), dorsal lateral prefrontal cortex (dlPFC) and posterior parietal cortex (pPC), selects a choice based on the subjective values. The mechanisms of how value accumulation interacts with subjective valuation to make a choice, and how brain regions communicate during decision making are undetermined. We developed and performed an EEG experiment that parametrically manipulated the probability of preferring delayed larger rewards. A computational model equipped with time and reward information transformation, selective attention, and stochastic value accumulation mechanisms was constructed and fit to choice and response time data using a hierarchical Bayesian approach.

Phase-based functional connectivity between putative dmFC and pPC was found to be associated with stimulus processing and to resemble the reconstructed accumulation dynamics from the best performing computational model across experimental conditions. By combining computational modeling and phase-based functional connectivity, our results suggest an association between value accumulation, choice competition, and frontoparietal connectivity in intertemporal choice.

**Author summary:** Intertemporal choice is a prominent experimental assay for impulsivity. Behavior in the task involves several cognitive functions including valuation, action selection and self-control. It is unknown how these different functions are temporally implemented during the course of decision making. In the current study, we combined formal computational models of intertemporal choice with a phase-based EEG measure of activity across brain regions to show that functional connectivity between dmFC and pPC reflects cognitive mechanisms of both visual stimulus processing and choice value accumulation. The result supports the notion that dynamic interaction between frontopatietal regions instantiates the critical value accumulation process in intertemporal choice.

## Introduction

The ability to make future-oriented decisions is a key component of long-term well-being [1–3]. Often, securing the largest cumulative reward requires forgoing immediate gratification, and holding out for a long term reward. A decision that requires an individual to choose among two or more rewards occurring at different time points is referred to as an intertemporal choice. Neuroimaging studies have identified a number of brain regions involved in intertemporal decision making [4–7], and scoped a general framework for understanding the functions implemented by these neural systems [8–10]. According to this framework, an intertemporal choice requires the completion of a subjective value representation, and a choice selection processes [4–6, 9, 11, 12]. To construct a subjective value representation, the information about the monetary value (i.e., reward) of a choice alternative and the amount of time (i.e., delay) associated with it must be fused together. This subjective value representation process is now widely acknowledged to be closely associated with neural processes in the ventral striatum and the ventromedial prefrontal cortex (vmPFC) across different experimental tasks [7, 8,12].

The process of choosing among intertemporal choice alternatives creates a type of competition for choice selection, and consideration of different aspects of the alternatives (e.g., reward and delay) creates different instantaneous subjective values, which must be compared. We have argued that value comparison occurs by accumulating stochastic evidence of the relative values of the choice options [13,14]. This choice value accumulation process has been associated with a large brain network, including dorsal medial frontal cortex (dmFC), dorsolateral prefrontal cortex (dlPFC) and posterior parietal cortex (pPC) [12,15,16]. Disrupting brain activity using transcranial magnetic stimulation (TMS) to bilaterally inhibit prefrontal cortex and right pPC increases impulsive behavior [11,17], implicating these brain regions in the consideration of future rewards. Harris et al. [18] exploited the temporal dynamics of intertemporal choice with EEG and found two separate roles of dlPFC: top-down attention filtering at early decision periods, and value modulation at late decision periods. Although there is extensive research and general consensus about the subjective valuation functions of vmPFC and ventral striatum, there is less consensus about the functional roles of dlPFC, dmFC and pPC because most previous studies have focused on only one specific region or mechanism. Such focus can be productive for localizing a region’s functional role, but it leaves open the question of how neural interaction between fronto-parietal regions contributes to decision-making. At least two important questions wait to be explored: (1) Do fronto-parietal regions assume separate or collaborative roles for making intertemporal choices? (2) What neural mechanisms most closely adhere to the value accumulation process?

To answer these questions, our current study uses the temporal resolution of EEG to probe the spatio-temporal covariates of cognitive mechanisms during intertemporal choice tasks. We first built a computational cognitive model that posits multiple potential mechanisms for the value accumulation process. We then determined the most plausible representation for the value accumulation process by factorially fitting different combinations of the model mechanisms to choice response time data hierarchically across subjects. Once the best mechanisms had been identified, we used the model to reconstruct the temporal dynamics of the value accumulation process, and associated these model dynamics with inter-regional phase-based functional connectivity among putative dmFC, pPC and dlPFC sources estimated from EEG data. Our results suggest that the functional connectivity between dmFC and pPC is a signature of the value accumulation process. Specifically, the dmFC-pPC functional connectivity, measured by theta-band phase clustering patterns in EEG data, is positively associated with the competitive dynamics of the value accumulation process. This finding enhances our understanding about the spatio-temporal neural correlates of intertemporal choice.

## Results

Subjects completed a pretest session and an EEG session of the intertemporal choice task. The purpose of the pretest session was to execute better control over subjects’ behaviors and to adequately decide choice offers for the EEG session. Specifically, we estimated two decision parameters for each subject according to a staircase procedure in the pretest session, such that we could tempt each subject to choose the delayed option with approximate probabilities of 0.1, 0.3, 0.5, 0.7, or 0.9 in the EEG session. We refer to such expected probabilities of choosing delayed options as experimental *P*_*D*_ conditions throughout the text.

In every trial of the EEG session, a delay *t* was randomly selected from a range of 15 to 45 days. We then calculated and offered an amount *r* that would give *P*_*D*_ of 0.1, 0.3, 0.5, 0.7, or 0.9; given the estimated parameters for the subject from the pretest session and the selected delay for the trial (see Methods section for details). Fig 1a shows how different *P*_*D*_ conditions were constructed (left) based on how each subject combined reward and delay information (right). For each subject, the combined rewards and delays constitute five discounted values (*V*_*D*_) of the delayed option, illustrated as five different colors on the plot. Then the *V*_*D*_s are mapped onto the five targeted *P*_*D*_ values, according to a sigmoid transform function. Fig 1b shows the temporal structure of stimulus presentation. Delays (*t*) were presented first for 1000 milliseconds. The amount information (*r*) was then shown and kept on screen until a choice was made or for a maximum of 4000 milliseconds. The immediate reward was the same ($10) on every trial, but it was only instructed to the subjects before the EEG session, instead of being presented visually. EEG data were collected during the EEG session.

**Fig 1.**
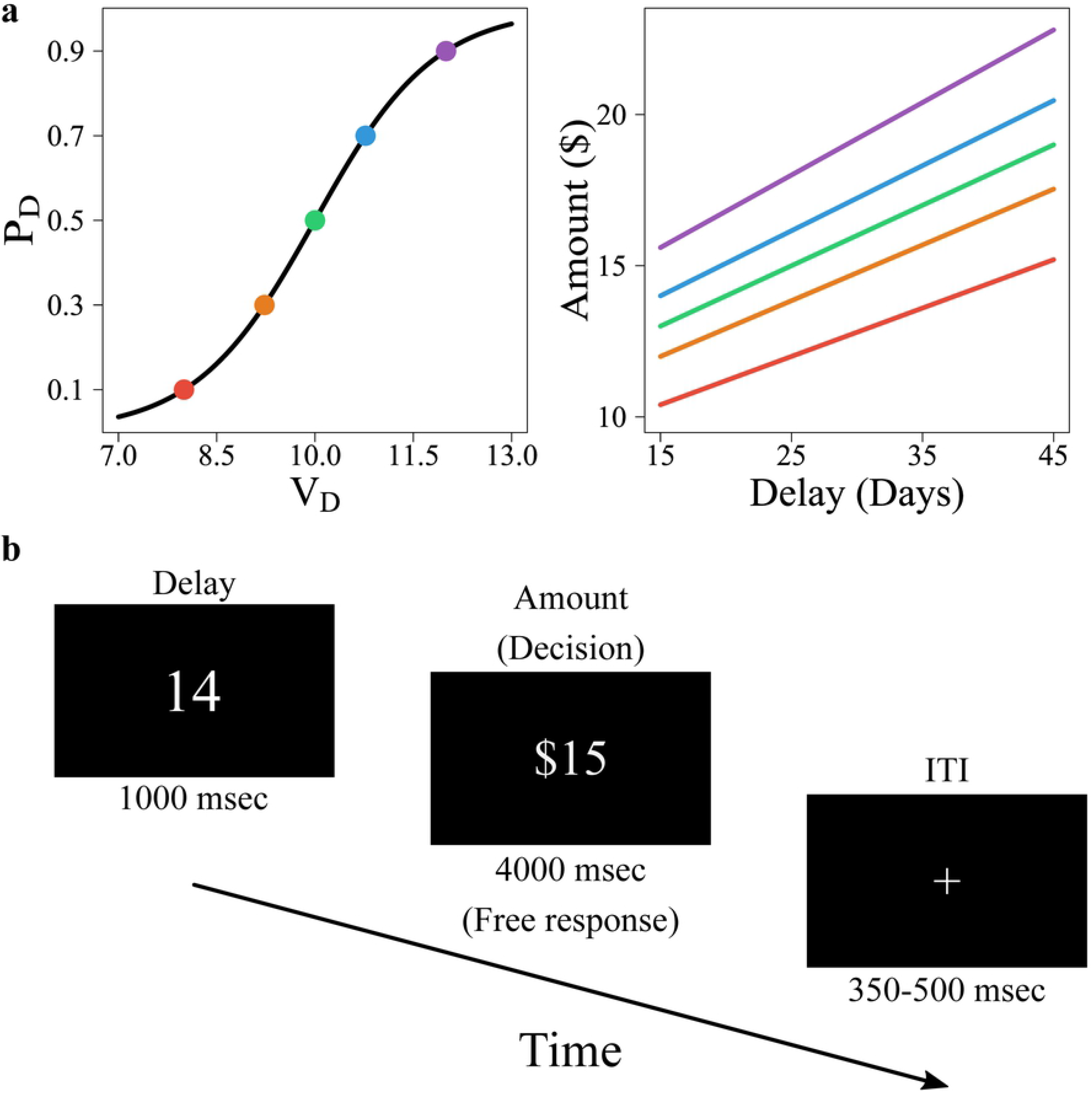
Experiment Design. (a) Discounted values of the delayed reward (*V*_*D*_) corresponded to one of five expected choice probabilities (*P*_*D*_) for each subject. Delays between 15-45 days were combined with different reward amounts (right panel) to map varying delayed rewards to targeted choice probabilities (left panel). (b) Within-trial sequential presentation of delay and amount information. Delays were presented first, followed by amounts. A jittered intertrial interval of several hundred milliseconds separated each trial.

### Behavioral data

To confirm that subjects’ behavior was consistent with the experimental manipulation, we first calculated the *empirical P_D_* as the proportion of delayed choice responses for each subject in each experimental *P*_*D*_ condition. Fig 2a illustrates that the empirical *P*_*D*_ (*y*-axis) during the EEG session closely matches the experimental *P*_*D*_ (*x*-axis) constructed from the pretest session. The solid bordered points correspond to the average empirical *P*_*D*_ across subjects, and they fall closely onto the diagonal line, suggesting that our experimental stimuli produced patterns of choice response we intended, despite some individual differences. To test that, we performed a 23 (subjects) *×* 5 (experimental *P*_*D*_ conditions) mixed effect logistic regression analysis with the *P*_*D*_ condition as a repeated measure and subject as a random effect. We found a significant effect of the experimental *P*_*D*_ condition, and the estimated probability of choosing the delayed option is 0.089, 0.287, 0.509, 0.759, and 0.869 for conditions *P*_*D*_ = 0.1, 0.3, 0.5, 0.7, and 0.9, respectively. The match between the experimental *P*_*D*_ and empirical *P*_*D*_ was thus confirmed.

**Fig 2.**
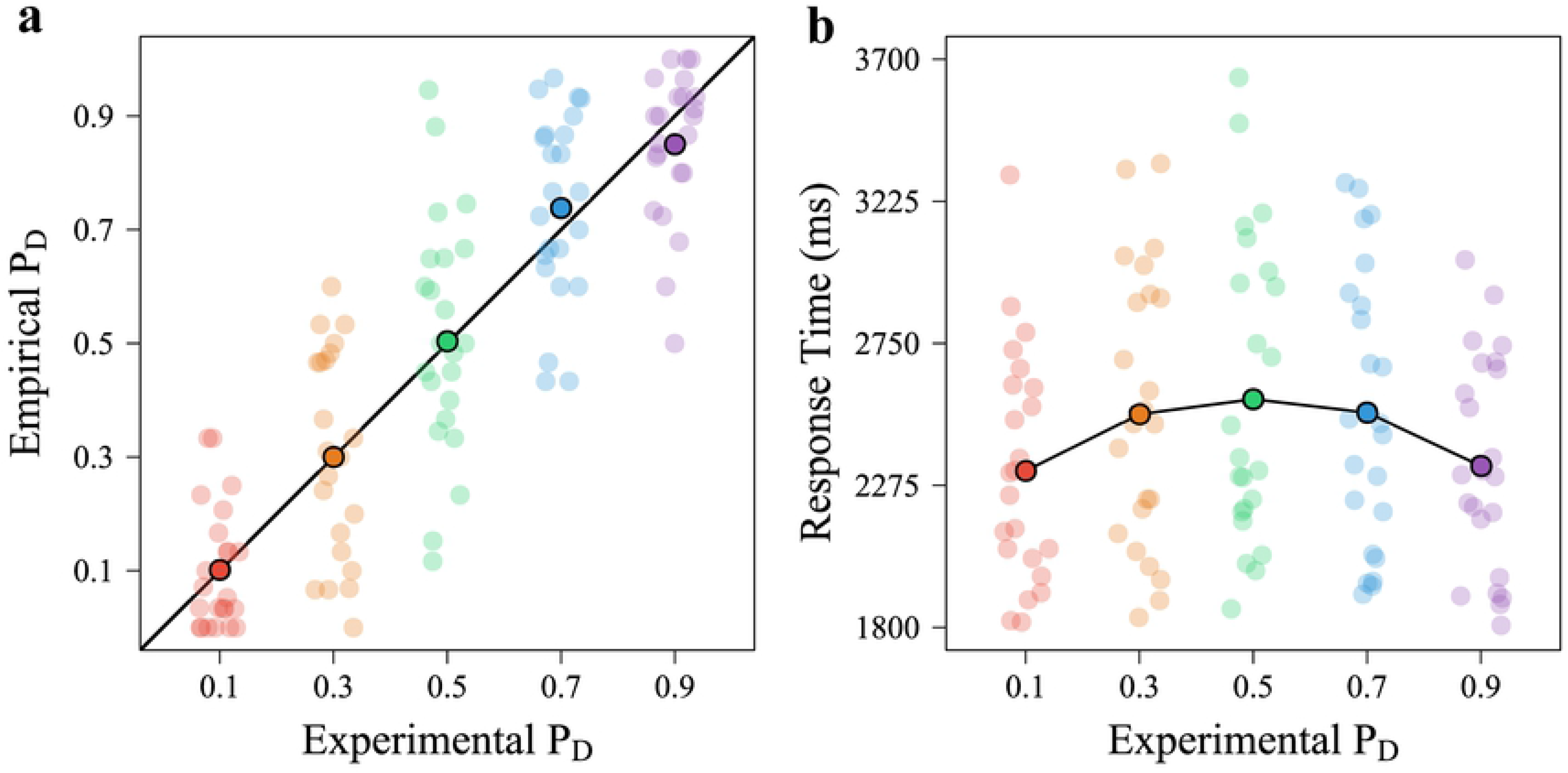
Behavioral Data Results. (a) The values of empirical *P*_*D*_ (actual proportion of making delayed choices) are consistent with the experimental *P*_*D*_ (theoretical probability of choosing delayed options). (b) Response times are shown against experimental *P*_*D*_. In both panels, the solid points correspond to the average value across subjects, whereas the transparent points correspond to the individual value for each subject. The black diagonal line in panel (a) illustrates the ideal equivalence between the empirical and experimental *P*_*D*_s.

Fig 2b depicts the mean response times for each subject (denoted as semi-transparent dots) as a function of the experimental *P*_*D*_. The average response time across subjects (denoted as the solid dots) shows a symmetric pattern where the average response time is longest at the *P*_*D*_ = 0.5 condition. To examine the symmetric relation between response times and *P*_*D*_ conditions, a mixed-effect regression model with orthogonal polynomial terms was performed, where the *P*_*D*_ condition was treated as a fixed effect and subject was treated as a random effect. The mixed-effect regression model suggested a significant effect of the quadratic experimental *P*_*D*_ on the response time (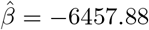, *t*(3999.01) = *−*11.396, *p* = 2.12 *×* 10^−16^) but did not yield significant results for the first-order term of *P*_*D*_ (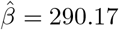, *t*(3999.01) = 0.512, *p* = 0.609). We also compared the quadratic model with a linear model in which the quadratic term was removed, and we found that the quadratic model generated both lower AIC and BIC values (AIC: 62524.41 and BIC: 62555.91) compared to the linear model (AIC: 62650.22, BIC: 62675.42). Such a comparison result indicated that the quadratic model better captured the response time data. Taken together, the behavioral choice probability and response time results corroborate that our experimental manipulation worked as intended.

### Behavioral modeling

In addition to analyses on manifest variables, we fit a computational model to the behavioral data to gain insight into the underlying within-trial temporal dynamics of intertemporal choice [15]. The model we adopted assumed a trade-off mechanism between features of rewards and time delays in forming the representation of choice alternatives [19]. Previous studies have shown that such attribute-wise models could mimic various delay discounting curves in terms of subjective valuation [15] and that these models explain choice probabilities and response times as well or better than hyperbolic discounting models do [20–22]. Depending on specific experimental designs, attribute-wise models can provide more information on temporal dynamics when different attributes were presented sequentially (as in the current experimental design). Similarly, attribute-wise models can be convenient to explain eye tracking data where the process tracing data reveal which attribute is being processed [22]. Fig 3a shows a graphical diagram of how the features of the stimuli (*r*_*I*_ , *r*_*D*_, *t*_*I*_ , *t*_*D*_) are mapped into a choice variable (*O*) from several model mechanisms, such as a transformation of time and reward values, moment-by-moment selective attention of attribute information, and lateral inhibition during the value accumulation process. The model takes its inspiration from decision field theory and its variants [21,23–25], and the leaky competing accumulator (LCA) model [26,27].

**Fig 3.**
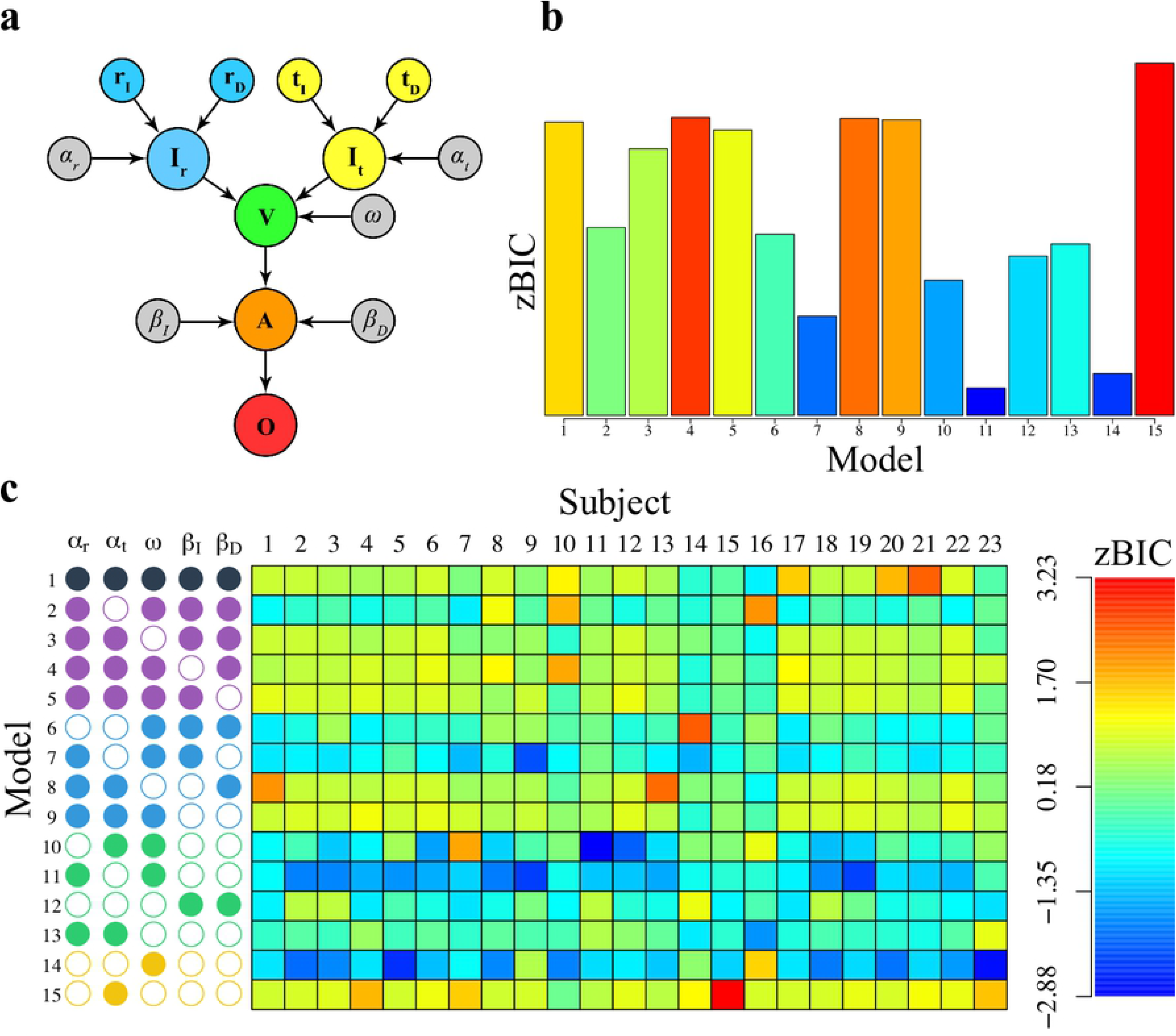
Computational Cognitive Model of Intertemporal Choice. (a) The model takes the objective value of rewards (i.e., *r*_*I*_ and *r*_*D*_; blue nodes) and time delays (i.e., *t*_*I*_ and *t*_*D*_; yellow nodes), and converts these inputs to subjective representations (i.e., *I*_*r*_ and *I*_*t*_) through transformation parameters *α*_*r*_ and *α*_*t*_, respectively. Features are selected at each moment with the parameter *ω*, creating an integrated subjective value of each alternative (i.e., the green node). Deliberation among the immediate and delayed alternatives is modulated by lateral inhibition parameters *β*_*I*_ and *β*_*D*_ (i.e., the orange node). Once an accumulator reaches a threshold amount of preference (*θ*), a decision is made corresponding to the winning accumulator (i.e., the red node). (b) Model fits from each model in (b), aggregated across subjects. (c) Model fitting results in terms of a z-transformed BIC statistic for each model configuration (by row) and each subject (by column). Each model variant is described on the left panel by a set of empty or filled circles, where empty circles indicate that a parameter was free to vary, whereas filled nodes indicate that a parameter was fixed. The model structures are grouped by their number of free parameters: black (3), purple (4), blue (5), green (6) and yellow (7). For the zBIC, lower values (blue) suggest better model performance.

In Fig 3a, the computational model converts information about the rewards (blue nodes) and time delays (yellow nodes) for both immediate (I) and delayed (D) options into subjective values. For this transformation, we assumed a power transformation [21]:

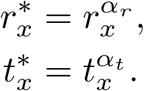

The power transformation is intended to represent how subjects encode stimulus values, where the reward *r*_*x*_ and delay *t*_*x*_ (*x* = *I* or *D*) information are transformed into subjective representations 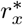 and 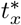 through the parameters *α*_*r*_ and *α*_*t*_, respectively.

The reward and time delay can be viewed as two separate feature dimensions that are selectively attended to at each moment in time *t*. Let *w*(*t*) denote the feature dimension that is being attended at the moment *t*. For simplicity, we assume that *w*(*t*) follows a Bernoulli distribution over time, such that

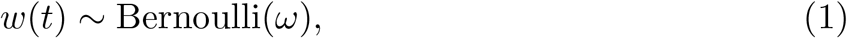

where *ω* is an attention parameter. We arbitrarily specified the values of *w*(*t*) such that when *w*(*t*) = 0, attention was directed toward the time information, whereas when *w*(*t*) = 1, attention was directed toward the reward information. Hence, estimates of the *ω* parameter that are large (i.e., near one) indicate greater attention to the reward information.

By selecting different featural dimensions to compare the immediate and delayed options over time, an integrated representation of the subjective values of each alternative (*V*_*I*_ and *V*_*D*_) can be constructed as:

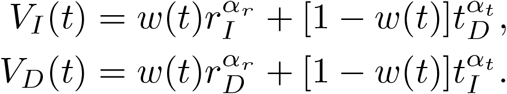

In Fig 3a, this isolated subjective valuation is represented as the green node. As shown in Fig 1, the time and reward information were not presented simultaneously, and so for the first 1000 milliseconds the time information could have been processed independently from the reward information (i.e. *w*(*t*) = 0). To account for this, we assumed that the subjective values were constructed in a “piecewise” manner, such that for the first 1000 milliseconds,

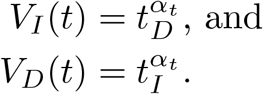

Hence, before the reward information is displayed (i.e., before one second), the subjective value of *V*_*I*_ and *V*_*D*_ only contains the encoded value of time delay. Once the reward information is presented (i.e., after one second), the subjective value oscillates between the encoded values of reward and time delay, with the frequency of attention determined by *ω* in Eq 1. Such a model formulation is strongly contingent on the specifics of the experimental design and the temporal resolution of the neural measurements. Given that we wished to connect the model’s temporal dynamics to EEG measurements with high temporal resolution, we determined that capturing this temporal contingency was crucial to successfully extracting a useful cognitive representation of subjective value accumulation.

Beyond the issue of subjective valuation is the issue of deliberation among the alternatives. Several studies suggest a key role for the effects of lateral inhibition and leakage on the temporal dynamics of decision making [15,16,21,23,26–31]. The lateral inhibition mechanism creates mutual competition between the immediate and delayed options during the course of value accumulation. The leakage mechanisms allow for the passive loss of information in the value accumulation process. For example, in our previous work [15], we have shown that trial-to-trial variability in the lateral inhibition parameter covaries with trial-to-trial measures of self control expressed in several of the integration brain regions discussed in the introduction (e.g., dmFC and dlPFC), yet were not linked to aspects of the valuation process (e.g., vmPFC).

Fig 3a illustrates how the subjective valuation node *V* is converted to the decision variable *A* with the mechanisms of lateral inhibition denoted *β*_*I*_ and *β*_*D*_, for the immediate (*I*) and delayed (*D*) options, respectively. *β*_*I*_ stands for the inhibition from the delayed option to the immediate option, and *β*_*D*_ stands for the inhibition from the immediate option to the delayed option. Although not illustrated in Fig 3a, the model also contains leakage mechanisms *λ*_*I*_ and *λ*_*D*_ to allow for the passive loss of information in the value accumulation process. Due to some parameter instabilities, in all model fits below, we do not estimate the leakage terms, but instead set them to *λ*_*I*_ = *λ*_*D*_ = 0.1 throughout. Finally, to allow for stochastic noise in the value accumulation process, we assumed the presence of Gaussian noise at each moment in time, such that

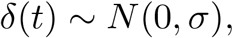

where *N* (*a, b*) denotes the normal distribution with mean *a* and standard deviation *b*. Unlike the leakage term, the noise term *σ* was estimated in all model variants.

Together, the value accumulation process including lateral inhibition, leakage and noise for each of immediate (*A*_*I*_) and delayed (*A*_*D*_) alternatives can be expressed as

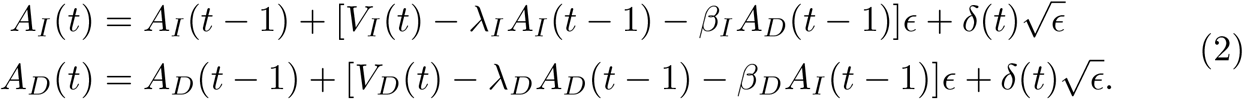

Although the model is actually a continuous diffusion process, Eq 2 is written as the recursive numerical approximation of the process in the Euler method [32]. The term *E* denotes a discrete time step in the accumulation process. We set *ϵ* = 0.1 in our approximation of the model’s dynamics.

There are two other dynamics of the model to discuss before finalizing the model specification, and both relate to boundaries that the accumulators *A*_*I*_ and *A*_*D*_ take. First, there is a threshold parameter *θ* that designates when enough value has accumulated for either option to trigger a response. On the presentation of the stimulus, both accumulators race to this threshold value *θ*, and a choice is made that corresponds to the accumulator that reaches the threshold first. Additionally, we specified that the accumulation process started at a fixed distance away from the threshold value *θ*. The starting point was fixed to be *θ*/5. Fig 3a illustrates that the accumulation process produces the response, designated as the red node *O*. For our purposes, we assumed a common threshold for both accumulators, and this parameter is estimated for every model variant we discuss below. Second, there is a lower boundary on the accumulation process, such that no accumulator can go below zero. This assumption follows the tradition of the Leaky Competing Accumulator model and is based off the principle that neurons cannot have negative firing patterns. To enforce this constraint, we specified that

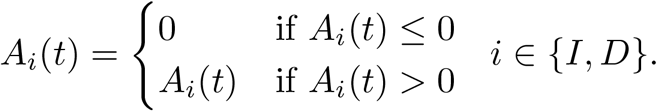

#### Model simulation

To help explain how the model works, Fig 4 illustrates the evolving value accumulation for the delayed (red) and immediate (blue) options under nine factorial combinations of delayed amount of reward (columns) and time delay (rows), compared to a fixed immediate $10 option. In each panel, multiple trajectories represent independently simulated trials to convey the across-trial variability with constant inputs of reward and time delays. As our experimental design first presents delay information, the first one-second of value accumulation only considers time (i.e., *ω*(*t*) = 0). As such, the accumulation during the first one-second period shows an advantage of the immediate option and the shape of the growth of preference is only affected by varied time delay (from top to bottom row), not by the amount of reward (from left to right columns). After the one-second period, the rate of value accumulation for the options accelerate based on a combination of both reward and delay, which can be seen by inspecting both rows and columns. Both alternatives compete to arrive at the threshold value first, at which time a corresponding response is triggered.

**Fig 4.**
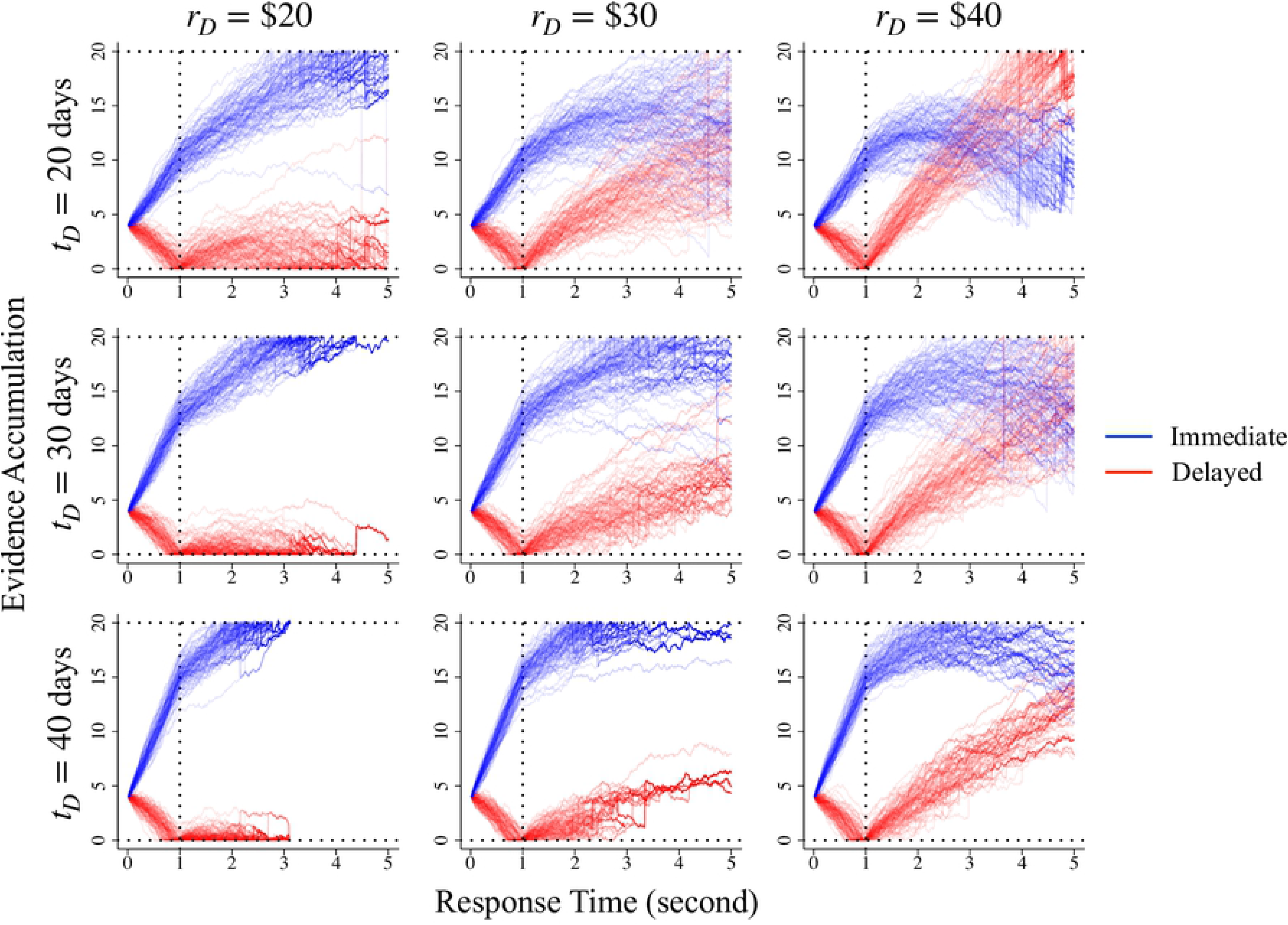
The Effects of Reward and Delay on Accumulation Dynamics. Simulated accumulation trajectories from the model under a factorial combination of reward (columns) and time delays (rows) for the delayed option, where the immediate option was held constant at $10 now. Each combination of reward and time delay was simulated 100 times, where the accumulation process starts as 0.2*θ* and ends once one accumulator reaches *θ*. In each panel, the immediate option is represented as blue trajectories, whereas the delayed option is represented in red. Other parameters were set to the following values: *α*_*r*_ = *α*_*t*_ = 0.7, *σ* = 1, *λ*_*I*_ = *λ*_*D*_ = 0.5, *β*_*I*_ = *β*_*D*_ = 0.5, *ω* = 0.9, *θ* = 20, *ϵ* = 0.01.

#### Model fitting and model selection

The computational model explains intertemporal choice behaviors as a series of processes that include value transformation, attention selection and value accumulation with leakage and lateral inhibition mechanisms. To determine which cognitive mechanisms are most important in terms of quantitatively accounting for the behavioral data, we adopted a Bayesian model selection approach. We first constructed different model configurations depending on which model parameters were freely estimated and which had fixed values. The left side of Fig 3c illustrates the fifteen different model variants we evaluated, where open or filled circles indicate that parameters *α*_*r*_, *α*_*t*_, *ω*, *β*_*I*_ , and *β*_*D*_ are either freely estimated or fixed for each individual, respectively. Each of the five parameters was set to a particular constant value when they were fixed and not freely estimated. Namely, we set *α*_*r*_ = 1, *α*_*t*_ = 1, *ω* = 0.9, *β*_*I*_ = 0, and *β*_*D*_ = 0. We chose to set *ω* to 0.9 because we suspected reward information would be prioritized asymmetrically because delay information was presented first and in isolation for the first second. Recall that for the first second, *ω* is set to zero for all models. In addition to these settings, we also freely estimated the accumulation threshold *θ*, moment-to-moment noise *σ*, and an additional non-decision time parameter *τ* for each model variant.

We fit each configuration to all behavioral data in a hierarchical Bayesian framework, meaning that each parameter just discussed was estimated for each subject, along with group-level parameters in the hierarchy (see Methods section for details of model parameter estimation). Once the fifteen model variants in Fig 3c were fit to data, we compared among the models by calculating a z-transformed Bayesian information criterion (BIC; [33]) metric, which is based on the model’s overall fit to data, penalized by the model’s complexity, and standardized across the model variants. Fig 3c depicts the degree to which each model configuration (rows) fit the behavioral data for each subject (columns), where lower values of zBIC (i.e., blue colors) indicate better model performance. Despite considerable individual differences in the best performing model, Fig 3c shows that some model configurations fit consistently better than others. In particular, Models 7, 10, 11, and 14 performed the best within this set of models. To better evaluate model performance at the aggregate level, Fig 3b shows the zBIC scores aggregated across subjects for each model. The aggregated model performance result affirmed the relative performance superiority of Models 7, 10, 11, and 14, with Model 11 performing particularly well.

To ensure that Model 11 is the best performing model among all tested model configurations, we further compared the best five models in Table 1 in terms of both log likelihood values and zBIC values, and we found that Model 11 and Model 14 fit almost equally well. Model 14 showed a slight advantage in log likelihood values but Model 11 had the lowest zBIC. To incorporate the models’ fits to individual subjects when evaluating the models, we ranked the zBIC values across model configurations for each subject, and counted the number of subjects having the lowest zBIC values for each model configuration. We also summed up the zBIC rank across subjects for each model to account for all the subjects. In this metric, having a lower summed rank indicates better performance by the model. Table 1 shows that Model 14 outperformed any other model for 9 out of 23 subjects, but it did struggle to fit certain subjects, such as subject 9, 14, and 16 (as suggested in Fig 3c). Model 11 worked best for 7 out of 23 subjects, and it had the lowest sum of zBIC rank. Model 11 also had a satisfactory fit to all the subjects according to Fig 3c. Together, we selected Model 11 as containing the most plausible representation mechanisms to explain our behavioral data. Hence, the model performed best when the following parameters were free to vary across subjects: *α*_*t*_, *β*_*I*_ , and *β*_*D*_. To get a sense of the optimal parameter values, we computed the maximum a posteriori (MAP) values from estimated posterior distributions for each parameter from both Model 11 (S1 Table) and Model 14 (S2 Table).

**Table 1.** Comparison of the Best Five Models. The log likelihood (or zBIC) values are computed by averaging log likelihood (or zBIC) values across subjects. “Count of lowest zBIC” is obtained by counting the number of subjects having the lowest zBIC given a certain model variant. “Sum of zBIC rank” sums up the individual rank of zBIC across subjects given a model variant.

| Model | Log Likelihood | zBIC | Count of lowest zBIC | Sum of zBIC rank |
| --- | --- | --- | --- | --- |
| 11 | −215.805 | −0.586 | 7 | 55 |
| 14 | −214.196 | −0.551 | 9 | 77 |
| 7 | −225.208 | 0.091 | 3 | 97 |
| 10 | −224.925 | 0.483 | 3 | 146 |
| 12 | −228.872 | 0.563 | 0 | 134 |

Note: The first column serves as the index for each model configuration, corresponding to the model index in Fig 3.

Although the model evaluation process assessed model performance in a relative sense (i.e., performance across models), it is also important to assess performance in an absolute sense. To do this, we evaluated how well the model fit the empirical data by simulating behavioral measures (i.e., choice response time) from the model, under the best parameter estimates obtained for each subject, 10 times for every trial and every subject. We inspected the simulation results by comparing the simulated behavioral data with the experimental data. Fig 5a illustrates the accuracy of Model 11’s (i.e., the best performing model) fit to the behavioral measurements. In each panel, summaries of simulated data from Model 11 are shown against the same summaries of the experimental data. The top panel shows the mean response times, whereas the bottom panel shows the probability of choosing the delayed option, for each subject and condition combination. Fig 5a shows that the majority of points fall on the diagonal line in the plot, suggesting close agreement of the model’s predictions and the empirical data. As another way to compare the simulated data against the experimental data for Model 11, we also plotted the choice and response time joint distributions across *P*_*D*_ conditions in S3 Fig, which again confirmed that the simulated data match experimental data. As a comparison, we also inspected Model 14, by comparing the simulated data and experimental data in terms of behavioral summaries, choice and response time distributions across *P*_*D*_ conditions (S4 Fig). Given the similar likelihood values of Model 11 and Model 14, it is not surprising that Model 11 (the best performing one) shows similar performance compared with Model 14 (the second best performing one), when evaluating aggregate model fit.

**Fig 5.**
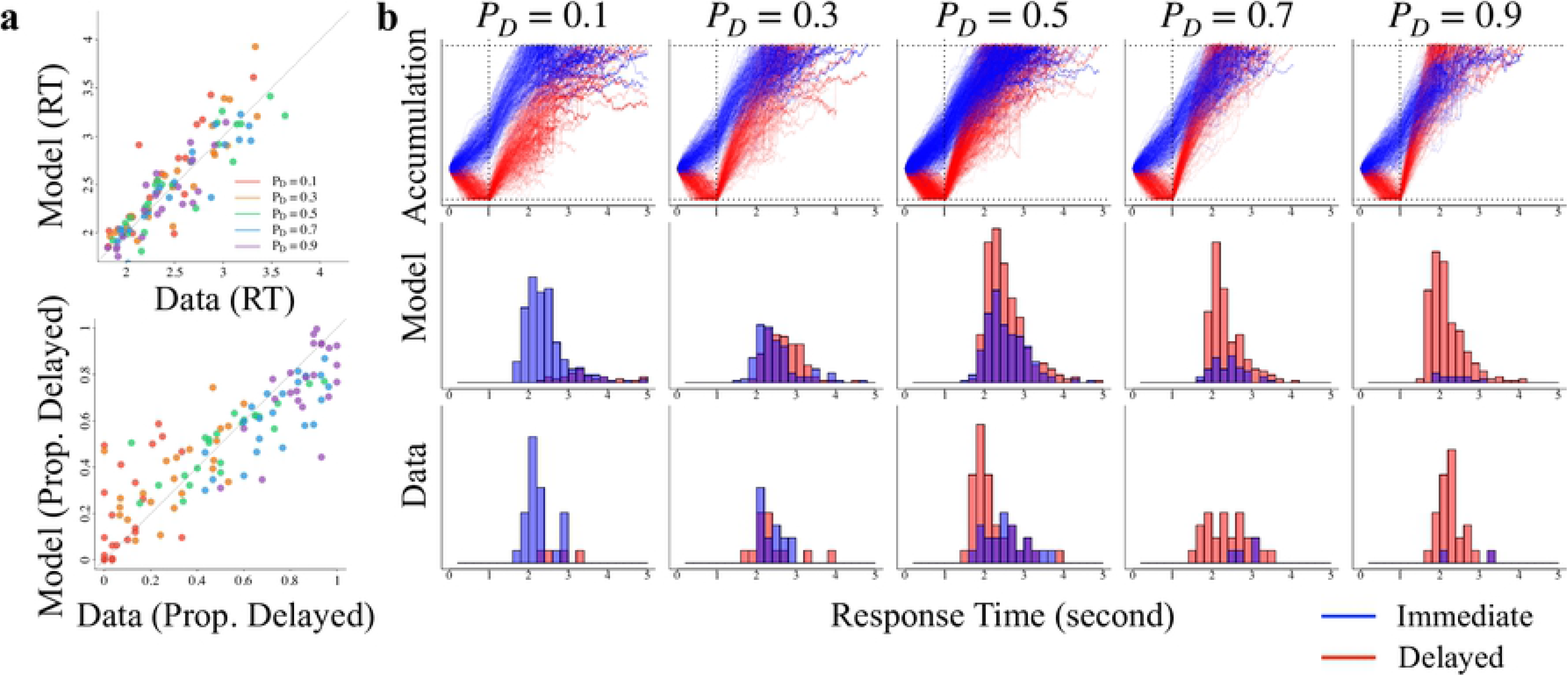
Model 11’s Fit to Data and Value Accumulation Representation. (a) The top panel shows the model’s predicted response time (RT) are shown against the experimental RT for each subject and condition, whereas the bottom panel shows the model’s predicted choice probability (for the delayed option) against the experimental data. (b) The columns show aspects associated with the five *P*_*D*_ conditions, whereas the rows correspond to simulated value accumulation dynamics (top), predicted choice RT distributions from the model (middle), and empirical choice RT distributions (bottom). In all panels, red colors correspond to delayed options, whereas blue colors correspond to immediate options. In panels in the middle and bottom rows, choice probabilities can be inferred by assessing the relative heights of the two histograms.

#### Value accumulation dynamics reconstructed from the behavioral model

The purpose of the model selection process was to identify the most plausible representation of value accumulation to explain the pattern of choices and response times across conditions. Our ultimate goal was to use this most plausible representation to interpret the temporal dynamics in our EEG data. Fig 5b provides an example of the model’s accumulation dynamics, and how well they fit to an individual subject. The top row shows the time course (0 - 5 seconds) of the modeled accumulation process for each of delayed (red) and immediate (blue) accumulators, where each line indicates one simulated trial. Across the five *P*_*D*_ conditions, the delayed and immediate accumulators race toward a common threshold. In each condition, the initial delay information causes some initial preference for the immediate option. However, once the reward information is presented (i.e., after one second), the delayed option gains preference, and the rate of this gain varies by condition, where the *P*_*D*_ = 0.1 condition is slowest, and the *P*_*D*_ = 0.9 is fastest. This rate differential produces a dynamic that results in the delayed option being strongly preferred in the *P*_*D*_ = 0.9. The impact of such a condition-wise pattern on the predicted behavioral output is shown in the middle row of the simulated data: a lower proportion of delayed choices are made in the *P*_*D*_ = 0.1 condition, a higher proportion is made in the *P*_*D*_ = 0.9 condition, and a approximately equal proportion of delayed and immediate choices are made in the *P*_*D*_ = 0.5 condition. These predictions closely match the shape of the choice RT distributions from the experiment, which are shown in the bottom row.

We applied a quantitative measure - absolute balance of evidence (ABOE; [34]) that summarizes accumulation dynamics for each experimental *P*_*D*_ condition. Specifically, we wanted to know how much competition occurred between the two alternatives across the conditions. In the model, competition reflects the degree to which each alternative is being considered, relative to the other alternative. Here, the difference between the evidence values of each accumulator is used to reflect competition: larger differences suggest less competition, whereas smaller values suggest more competition. Fig 6a conceptually illustrates the calculation of the ABOE measure with two single-trial examples of value accumulation. The left panel shows a situation in which competition is low, whereas the right panel shows a situation in which competition is high. For the two single trials, ABOE is conceptualized as the distance between the evidence of the two accumulators at the time a choice is made. In the case of low competition, the immediate accumulator keeps growing before reaching the threshold, while the delayed accumulator climbs with a lower speed and is unable to catch up to the immediate accumulator. As a comparison, in the case of high competition, the delayed accumulator rises after one second at a fast rate, gradually catching up with the immediate accumulator and reaching the threshold first. Across panels, the distance between the accumulators was larger in the low competition scenario relative to the high competition scenario.

**Fig 6.**
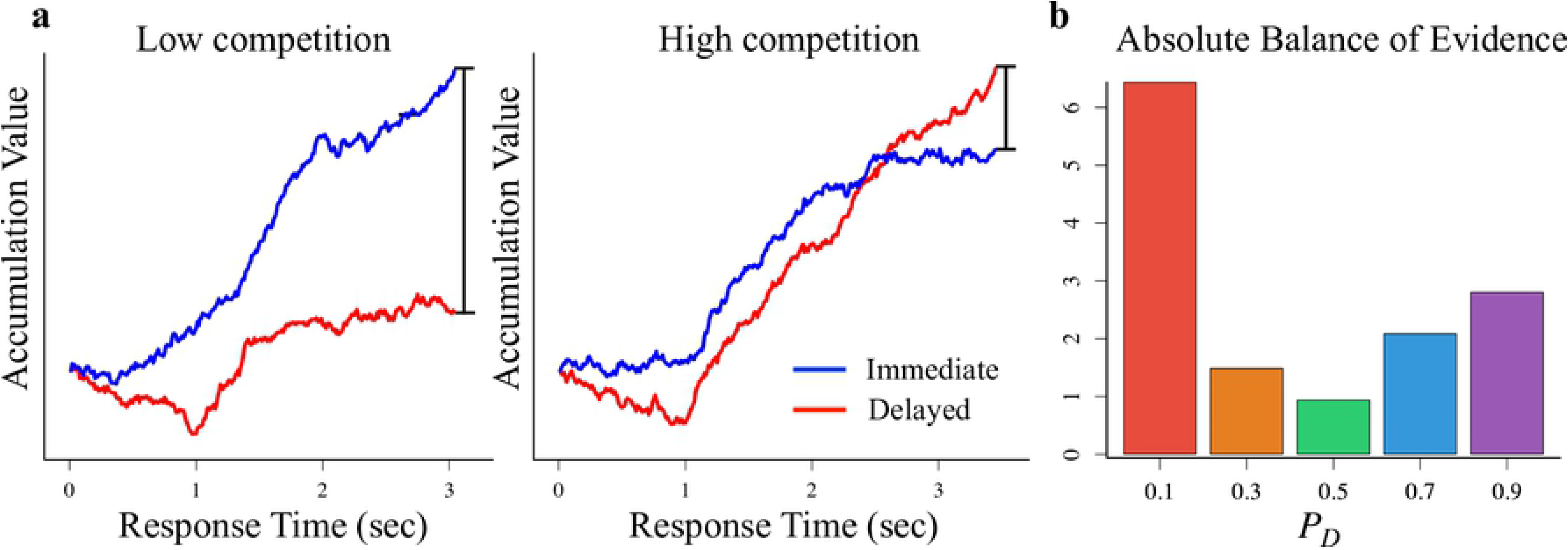
The Calculation of Absolute Balance of Evidence (ABOE). (a) Two single-trial examples of the value accumulation process with low (left) or high (right) competition between the two accumulators. For the two single trials, ABOE is conceptualized as the distance between the evidence of the two accumulators at the time a choice is made (denoted by the length of the line segments). (b) Resulting ABOE of the model simulations across the five *P*_*D*_ conditions.

After calculating the ABOE for each simulation, we averaged the ABOEs across conditions, such that

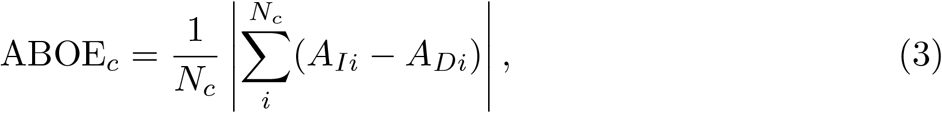

where ABOE_*c*_ refers to the absolute balance of evidence for condition *P*_*D*_ = *c*, *N*_*c*_ refers to the number of simulated trials in experimental condition *P*_*D*_ = *c*, and the *A*_*Ii*_ and *A*_*Di*_ refer to the instantaneous evidence of simulated trial *i* of immediate and delayed accumulators at the choice time, respectively. Fig 6b compares ABOE across five *P*_*D*_ conditions. The ABOE value was lowest at condition *P*_*D*_ = 0.5, and was highest at condition *P*_*D*_ = 0.1. Such a pattern of ABOE across *P*_*D*_ conditions suggests that choices were most difficult to make in the *P*_*D*_ = 0.5 condition, but were relatively easier in the *P*_*D*_ = 0.1 and *P*_*D*_ = 0.9 conditions. Taken together, the simulated accumulation processes depicted in Fig 5b and the summary measure ABOE in Fig 6b suggest a symmetric pattern from *P*_*D*_ = 0.1 to *P*_*D*_ = 0.9, where the accumulation process was the most difficult due to competition at the *P*_*D*_ = 0.5 condition.

Theoretically, by virtue of the ABOE measure, one could identify different degrees of value competition between the reward amount and time delay that was accumulated during the trial. The first second in the time course, however, does not contain any presentation of reward amount, and so the degree of value accumulation in this time period is only driven by the magnitude of time delay. This particular time window has to be accounted for separately when we summarize the value accumulation dynamics.

### Temporal dynamics of functional connectivity from EEG

#### Electrodes and ERPs

We analyzed the event-related potentials (ERPs) on several electrodes of EEG data after the extracted epochs were transformed to current source density (CSD) estimates (see Methods section). The ERPs were calculated based on a set of electrodes that were considered to correspond to dmFC, left/right pPC, and left/right dlPFC. Table 2 listed the different sets of electrodes in the analysis, the approximately corresponding electrodes in the International 10/20 system, and the respective putative brain areas. We included those electrodes in the ERPs analysis and the following ISPC analysis based off previous studies. For example, the putative electrode E6 that represents dmFC region corresponds to FCz electrode in 10/20 system [35,36], and the FCz electrode has been implicated with decision related functions in other domains [37,38]. The location of F3 and F4 have been localized to Brodmann area 46 [39], roughly corresponding with dlPFC [40]. The location of TP3 and TP4 have been localized to Brodmann area 40 (or the inferior parietal lobule) [39], which constitutes the lateral part of pPC [41].

**Table 2.**
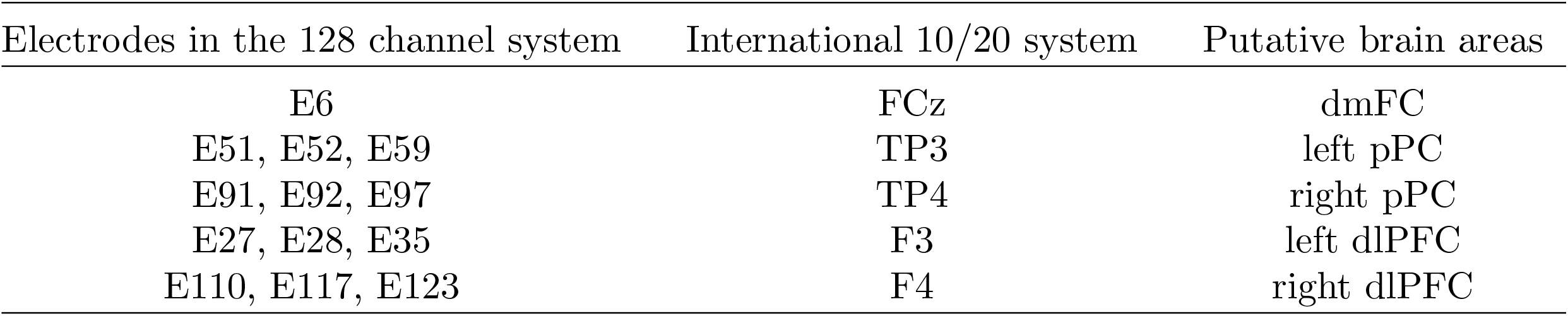
Details of the Electrodes Used in the ERPs and ISPC analyses.

Fig 7 illustrates the ERPs calculated based on these electrodes, representing brain activity within dmFC, bilateral dlPFC, and bilateral pPC, respectively. We separated trials based on *P*_*D*_ conditions to examine whether there is any difference in EEG amplitude associated with *P*_*D*_. We focused on two time windows (0-500ms and 1000-1500ms) each following the presentation of time delay and reward amount, due to the visual increase after each stimulus presentation. For each putative brain area, we ran a two-way mixed ANOVA with two fixed factors (*P*_*D*_ condition and time window) and subject as random factor. The analyses did not yield any significant difference in the EEG amplitude for dmFC, in terms of either *P*_*D*_ (*F*(4, 202) = 0.2004*, p* = 0.938) or time window (*F*(1, 202) = 3.2034*, p* = 0.075). The analysis performed on dlPFC also suggested no significant difference in EEG amplitude for either *P*_*D*_ (*F*(4, 202) = 0.5247, *p* = 0.718) or time window (*F*(1, 202) = 3.4534, *p* = 0.065). For pPC, we identified a significant difference between the two time windows (*F* (1, 202) = 41.6499, *p* < 0.0001), but no significant difference on *P*_*D*_ (*F*(4, 202) = 0.3405, *p* = 0.850). Next, we tested whether the EEG amplitude in each time window and each brain area was significantly different from the baseline level. For dmFC, the EEG amplitude in both time windows was significantly different from zero (0-500ms: *t*(22) = 2.9109, *p* = ,008; 1000-1500ms: *t*(22) = 3.1187*, p* = 0.005). For pPC, the EEG amplitude in time window 0-500ms was significantly different from zero (*t*(22) = 2.7463*, p* = 0.012). Other tests did not yield significant result (pPC 1000-1500ms: *t*(22) = 0.818*, p* = 0.422; dlPFC 0-500ms: *t*(22) = 0.4567*, p* = 0.652; dlPFC 1000-1500ms: *t*(22) = 1.157, *p* = 0.260)

**Fig 7.**
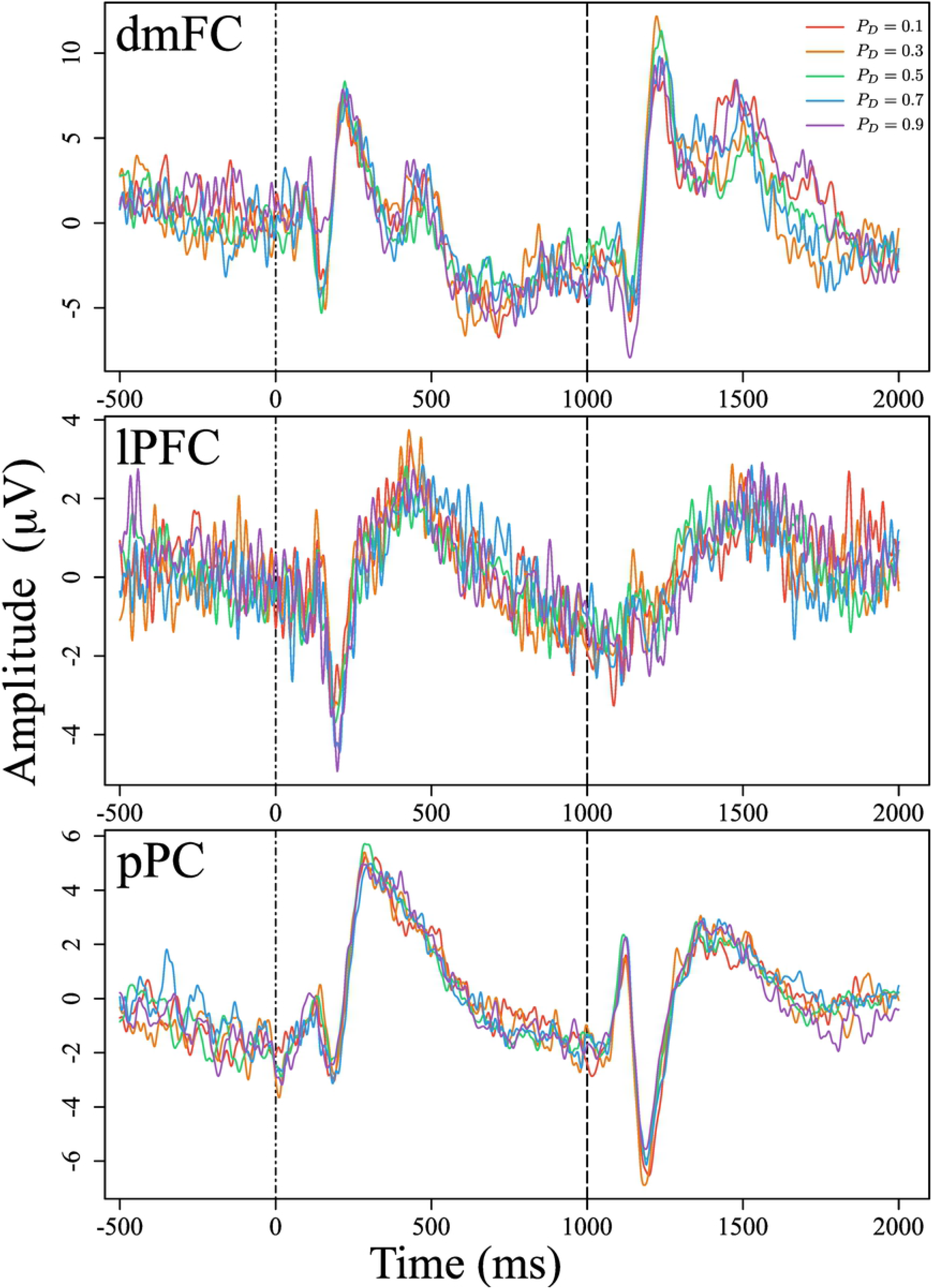
ERPs from Putative dmFC, dlPFC, and pPC for each *P*_*D*_ condition. ERPs were calculated on the preprocessed EEG data after CSD transformation to reduce volume conduction. Three panels from top to bottom show scalp electrical activities approximately located to dmFC, dlPFC, and pPC, respectively. Trials were aligned by the onset of time delay presentation, and the time course extends from −500 ms to 2000 ms. Two dashed lines indicate the presentation of time delay (0ms) and reward amount (1000ms). Trials were separated by *P*_*D*_ conditions for comparison.

### pISPC between putative dmFC, pPC, and dlPFC

We analyzed functional connectivity in the EEG data by calculating intersite phase clustering (ISPC). ISPC measures the extent to which phase clustering patterns are similar across different sites, therefore serving as a measure of the time-dependent connectivity strengths between putative brain regions [42]. The pISPC standardizes ISPC by calculating the percent change in ISPC with respect to the grand average ISPC over the time window of interest (see Methods section). We chose dmFC as the seed region, as we have previously found its association with value accumulation activity using fMRI [43]. The connectivity was established between dmFC and other regions previously identified important in intertemporal choice, including bilateral pPC and bilateral dlpFC. Each brain area was localized by a set of electrodes listed in Table 2.

Fig 8a shows the calculated pISPC result between putative dmFC and left pPC (left), and between putative dmFC and right pPC (right), as a function of time and frequency. In each panel, two clusters of increased pISPC emerge after trial onset and after one second in the trial time course, where the increase of pISPC after one second occurs at around 4 - 8 Hz for both dmFC-left pPC and dmFC-right pPC. The increase of pISPC after trial onset occurs at a wider frequency band: around 7 - 15 Hz for dmFC-left pPC and around 4 - 15 Hz for dmFC-right pPC. Note that in our experimental design, the time points zero and one second correspond to the presentation of delay and reward amount, respectively. Fig 8b shows the functional connectivity between dmFC and dlPFC based on the pISPC. In contrast with dmFC-pPC, the connectivity between dmFC and dlPFC did not yield the obviously increased pISPC. Although previous studies have identified gamma frequency band (30 - 90 Hz) in EEG activity as reflecting evidence accumulation mechanisms [44,45], the present study localized the informative frequency band to lower frequency, especially to theta band (4 - 8 Hz). Therefore, we focused subsequent analyses on theta band activity and tested for increases of pISPC following the stimulus presentation by analyzing pISPC within the time windows 0 - 0.3 s and 1 - 1.3 s (demarcated by black borders in Fig 8). Following the presentation of the time delay information, there was an increase of functional connectivity between dmFC and right pPC, but not between dmFC and left pPC (left: *t*(22) = 1.9829, *p* = 0.06; right: *t*(22) = 2.5135, *p* = 0.0198). Following the presentation of the reward information, there was an increase of functional connectivity between dmFC and bilateral pPC after the presentation of the reward amount (left: *t*(22) = 3.1714, *p* = 0.0044; right: *t*(22) = 3.0492, *p* = 0.0059). Similarly, we performed one-sample *t*-tests to investigate functional connectivity between dmFC and dlPFC. However, only the left dlPFC yielded a significant difference after the presentation of the time delay (*t*(22) = 2.2723, *p* = 0.0332), whereas other combinations did not show statistical differences neither after the presentation of the time delay (right: *t*(22) = 1.9237, *p* = 0.0674), nor after the presentation of the reward (left: *t*(22) = 1.8397, *p* = 0.0793, right: *t*(22) = 0.8363, *p* = 0.4120). Taken together, the functional connectivity between dmFC and dlPFC did not show obvious activation patterns compared to dmFC-pPC. Hence, we focused on the functional connectivity between dmFC and pPC for subsequent analyses.

**Fig 8.**
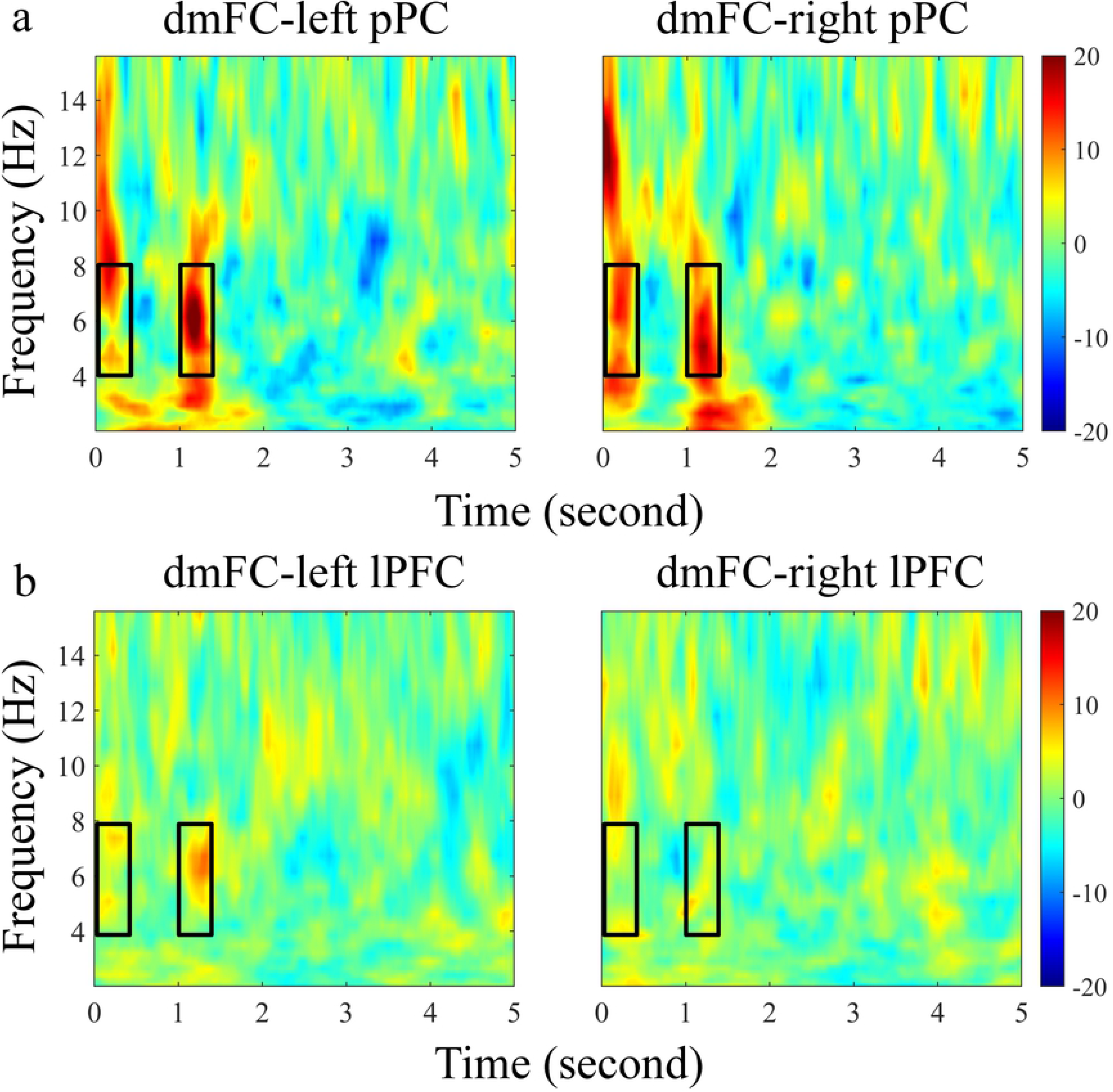
Functional Connectivity among Putative dmFC, pPC, and dlPFC. Each time-frequency plot shows how functional connectivity (pISPC) changes across the trial time course (0 - 5 seconds) and frequency values, where 0 seconds on the x-axis corresponds to the onset of delay information. Each plot averages all subjects and five *P*_*D*_ conditions. The black boxes indicate the selected time-frequency range (theta band; 0 - 0.3 s and 1 - 1.3 s) for performing associated statistical analyses.

### dmFC-pPC pISPC across *P*_*D*_ conditions

Given the evidence of increased functional connectivity at two time windows, we subsequently explored the possible mechanisms underlying this result. Because the functional connectivity results temporally coincide with the presentation of the delay and amount information, we suspected that the functional connectivity between the pPC and dmFC may simply reflect visual processing of stimulus information. To examine this possibility, we performed a repeated *t*-test between the two time windows for both the left and right pPC, but found no significant evidence of a statistical difference (left: *t*(22) = 1.4916, *p* = 0.15; right: *t*(22) = 0.6433, *p* = 0.6433), further suggesting that the connectivity may be related to visual stimulus processing. However, many previously established studies suggest that the time delay and reward amount are integrated into a subjective representation, and a subsequent value accumulation process determines the choice. The key question that our experiment is well poised to address is whether the dmFC-pPC connectivity can provide insight into the cognitive processes that underlie these choices.

To investigate whether connectivity is related to the cognitive aspects of our task, rather than just visual information, we tested whether neural communications between the two regions are mediated by the subjective valuation of the stimuli. To this end, we performed similar time-frequency analyses of the functional connectivity between putative dmFC and putative pPC, separated by each *P*_*D*_ condition. Fig 9a illustrates how temporal dynamics of functional connectivity at different frequency bands vary across five *P*_*D*_ conditions. As the increase of connectivity is mostly contained within the time course of 0 - 2 seconds, we omitted plotting other time ranges for visual clarity. We observed that the two clusters of increased pISPC exhibit varied magnitudes across *P*_*D*_ conditions, and this pattern appears symmetric across the five conditions: the magnitude increases from *P*_*D*_ = 0.1 to *P*_*D*_ = 0.5, and then decreases from *P*_*D*_ = 0.5 to *P*_*D*_ = 0.9, where the magnitude is highest at condition *P*_*D*_ = 0.5. This symmetric variation is precisely the pattern predicted by our ABOE measure of value accumulation dynamics, but this pattern is not predicted from the visual properties of the stimuli alone. Furthermore, the symmetric pattern of pISPC was found only after the presentation of the reward information, and not the presentation of the time information (i.e., the first aspect of the stimulus). Taken together, we conclude that the connectivity result is unlikely to be driven purely by stimulus processing interactions alone, and may instead be modulated by a value accumulation mechanism.

**Fig 9.**
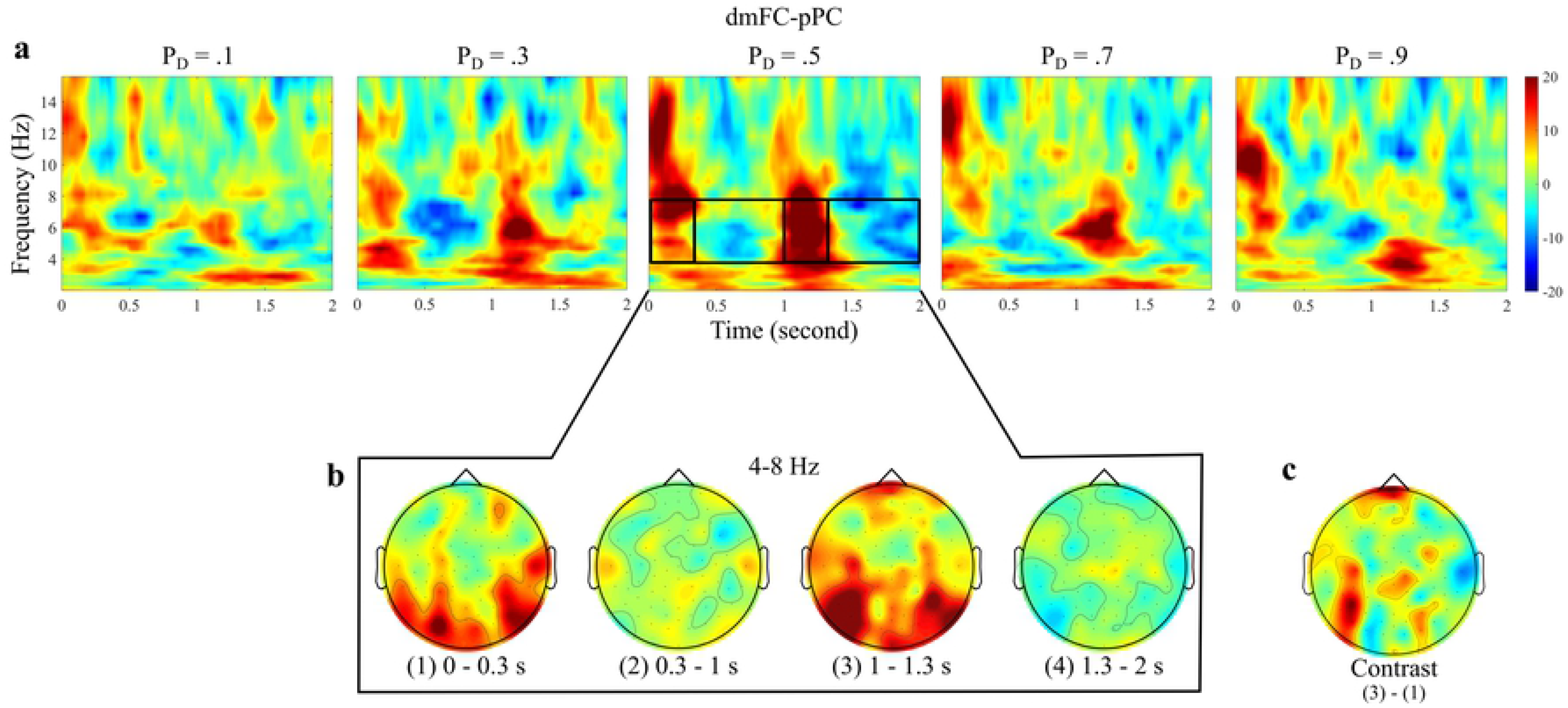
Functional Connectivity between Putative dmFC and Bilateral pPC across P_D_ Conditions. (a) Time-frequency analyses of pISPC between putative dmFC and pPC for each experimental *P*_*D*_ condition from 0.1 to 0.9, where zero seconds on the *x*-axis correspond to the onset of delay information. The time-frequency plots show the average connectivity values of dmFC-left pPC and dmFC-right pPC. (b) Topographical plots of functional connectivity between E6 (putative dmFC) and all other electrodes across four separate time windows within the frequency band 4 - 8 Hz at *P*_*D*_ = 0.5. (c) A contrast topographical plot showing the difference of ISPC values between the time window 3 (i.e. 1 - 1.3 seconds) and time window 1 (i.e. 0 - 0.3 seconds) in (b).

We further examined the topographical characteristics across time windows for the *P*_*D*_ = 0.5 condition, which had the largest pISPC between dmFC and pPC. Fig 9b shows topographical plots of the pISPC between the seed region E6 (putative dmFC) and all other scalp electrodes, as an indicator of the change of connectivity across space and time. The connectivity is stronger between E6 and parietal electrodes than between E6 and all other electrodes, indicating that frontoparietal functional connectivity dynamics are dominant compared with frontal connectivity dynamics. Such strong frontoparietal connectivity exists after both delay and reward information, but yields stronger activation following reward information. To better illustrate this effect, Fig 9c shows the contrast between the topographical connectivity values between 1 - 1.3 seconds and 0 - 0.3 seconds, where the difference is most pronounced in the left parietal areas. As such, although there is no statistically significant difference between the two time windows for the dmFC-pPC connectivity, the topographical plots suggest the existence of a difference when considering the large set of left posterior electrodes. This result further supports the notion that the increase in connectivity in the 1 - 1.3 second time window is not only simply about processing of visual information; rather, it may be associated with the competition between two value accumulators.

### Theta-band dmFC-pPC pISPC across *P*_*D*_ and across time delays

To closely examine the possibility that the subjective valuation process mediates functional connectivity, Fig 10a compares the pISPC across the five *P*_*D*_ conditions within theta frequency band of 4 - 8 Hz. The locally maximum pISPC results occur at around time 0 - 0.3 seconds and 1 - 1.3 seconds, where the height of each curve is modulated by experimental condition. Fig 10c shows the average of the pISPC result corresponding to the each presentation of delay and amount information. We performed a mixed-effect regression analysis of pISPC with the experimental *P*_*D*_ as the predictor variable and subject as the random variable. To observe any quadratic trends with *P*_*D*_, we included both first-order and second-order *P*_*D*_ terms in the regression model. We fit the regression model on both time windows, and the result indicated no significant quadratic association between *P*_*D*_ and pISPC at the time window 0 - 0.3 seconds (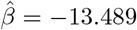, *t*(92) = *−*1.139, *p* = 0.2575), but a significant quadratic association between *P*_*D*_ and pISPC at the time window 1 - 1.3 seconds (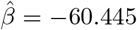, *t*(92) = −4.710, *p* = 8.74*e* − 06). We did not observe any linear association between *P*_*D*_ and pISPC on either time window. Therefore, consistent with the quadratic association between *P*_*D*_ and response time in Fig 2b, we observed a quadratic relation between the experimental *P*_*D*_ and pISPC on time window 1 - 1.3 s for dmFC-pPC connectivity. We also performed the same analyses based on separate dmFC-left pPC and dmFC-right pPC, and obtained the same pattern of results. Specificially, there were significant quadratic associations between *P*_*D*_ and pISPC at the time window 1 - 1.3 seconds (left: 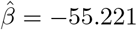, *t*(92) = *−*3.380, *p* = 0.00106; right: 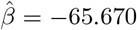, *t*(92) = *−*4.155, *p* = 7.29*e −* 05), but were not significant quadratic associations at the time window 0 - 0.3 seconds (left: 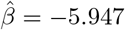, *t*(92) = *−*0.397, *p* = 0.6922; right: 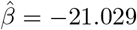, *t*(92) = *−*1.388, *p* = 0.1685).

**Fig 10.**
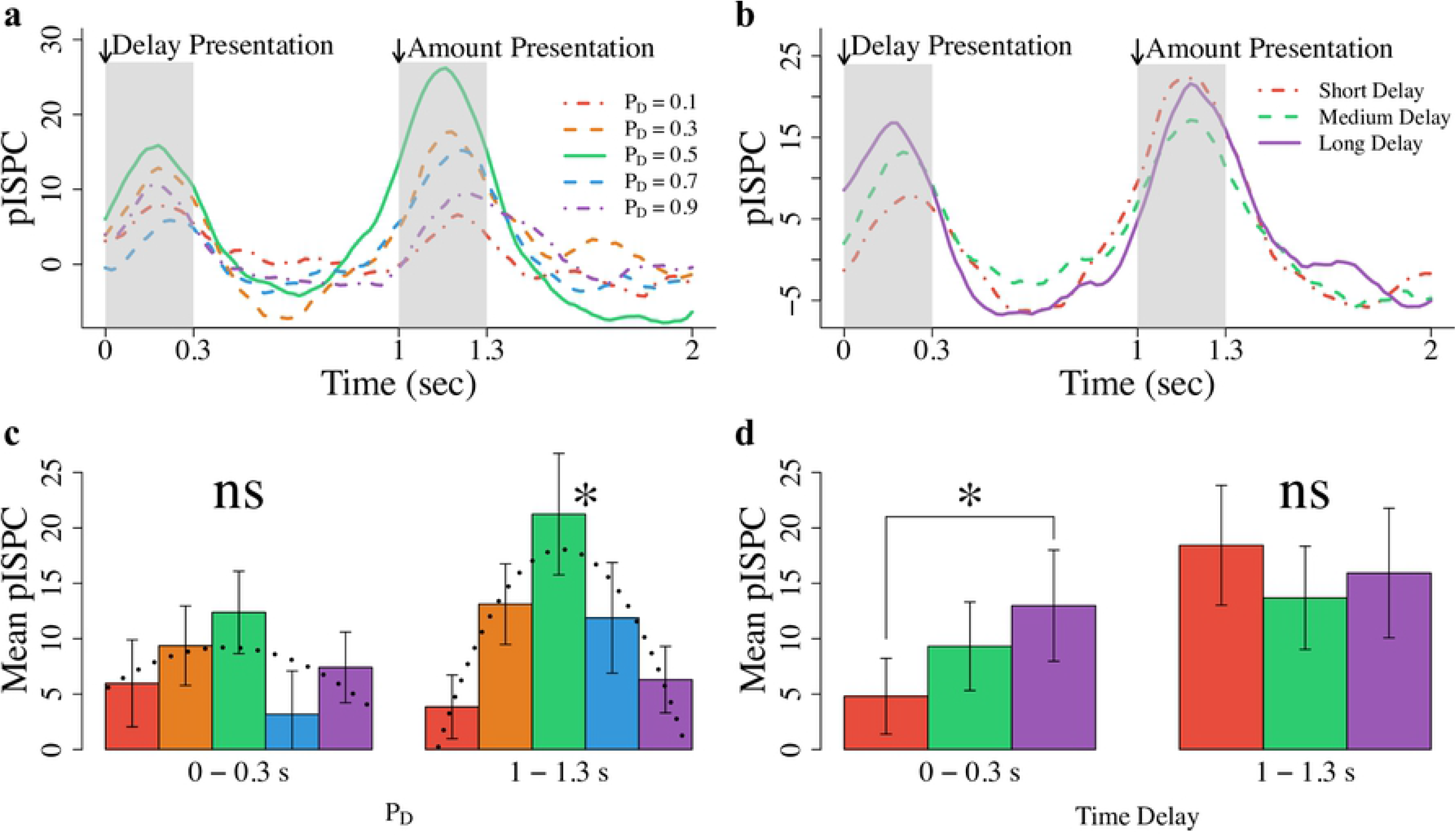
Comparison of the dmFC-pPC pISPC across P_D_ Conditions and Time Delays. (a) The pISPC values across time for each *P*_*D*_ condition (lines), where pISPC is calculated between dmFC and bilateral pPC. The two time windows shaded with gray denote the 0 - 0.3 second period (i.e., after delay presentation) and 1 - 1.3 second period (i.e., after amount presentation). (b) The pISPC values across time for each time delay condition (lines), where pISPC is calculated between dmFC and bilateral pPC. The three time delay conditions correspond to time delays of 15 - 25 days, 26 - 35 days, and 36 - 45 days. (c) The mean pISPC for each *P*_*D*_ condition within 0 - 0.3 second period and within 1 - 1.3 second period. The asterisk denotes a significant quadratic association at the *α* = 0.05 level, whereas “ns” denotes no significant quadratic association at the *α* = 0.05 level. The dotted lines indicate the fitted quadratic regression functions. (d) The mean pISPC for each time delay condition within 0 - 0.3 second period and within 1 - 1.3 second period. The asterisk denotes a significant pairwise difference at the *α* = 0.05 level, whereas “ns” denotes no significant difference across conditions at the *α* = 0.05 level. For both c and d, the error bars denote the *±*1 standard error where values below zero are not shown on the plot.

We observed that the average of pISPC varied across experimental *P*_*D*_ during time window 1 - 1.3 s, but not during 0 - 0.3 s. This result is consistent with our experimental manipulation, as subjects were only shown the time delay information at 0 - 0.3 s, and so we do not expect the role of *P*_*D*_ during this time period. On the contrary, because the time delay is the only information presented before 1 second, it is reasonable to consider whether the dmFC-pPC connectivity is modulated by the magnitude of time delays, similar to the observation that accumulation dynamics were only affected by time delay in Fig 4. To investigate this possibility, we binned the time delays across all subjects to three different groups: Short Delay (15 - 25 days), Medium Delay (26 - 35 days), and Long Delay (36 - 45 days) and compared the dmFC-pPC connectivity associated with each group. Fig 10b shows the pISPC measure corresponding to each of the delay groups across 0 - 2 seconds. Interestingly, we observed a clear separation of pISPC magnitude during 0 - 0.3 s, but not during 1 - 1.3 s. We computed the average value of pISPC within each time window for each delay group, illustrated in Fig 10d. We carried out a repeated measure ANOVA with the delay information as an independent variable for both time windows. The ANOVA model suggested a significant difference across three delay groups in the 0 - 0.3 second period (*F*(2, 44) = 3.457, *p* = 0.0403). The post-hoc analysis with Bonferroni’s correction further suggested a significant difference between Long vs. Short comparison (*t*(44) = 2.625, *p* = 0.0313), but non-significant results on other two comparisons (Medium vs. Short: *t*(44) = 1.447, *p* = 0.3264; Long vs. Medium: *t*(44) = 1.178, *p* = 0.4723). The same repeated measure ANOVA was performed for the 1 - 1.3 second period, but it did not yield any significant results (*F*(2, 44) = 0.588, *p* = 0.56). Together, we concluded that the magnitude of pISPC was modulated by the time delay in the 0 - 0.3 second period, with higher pISPC associated with longer time delay.

Combining the above findings during 0 - 0.3 s and during 1 - 1.3 s, we observed that the magnitude of pISPC is modulated by the time delay before the amount of reward is presented, and is afterwards modulated by the experimental *P*_*D*_, which represents the integral information of both time delay and amount of reward. Given that experimental *P*_*D*_ is closely associated with the amount of value competition between two choice alternatives, pISPC can be viewed as an indicator of general value accumulation, such that it tracks either the value of time delay when only time delay is available, or the amount of value competition when both time delay and amount of reward are available.

## Discussion

In this article, we have shown that fronto-parietal connectivity is an expression of valuation accumulation within an intertemporal choice task. We have also shown that a computational model in which subjective values are constructed through moment-to-moment sampling of feature dimensions exhibits patterns of choice competition that closely resembles the pattern of connectivity results across conditions of our experiment. In this section, we first discuss the relevance of the cognitive mechanism we identified from our behavioral data. We then discuss the connectivity results, and what they might suggest from a cognitive perspective. Finally, we discuss how our neural results can be more fully integrated into computational models in future work.

### Cognitive mechanisms of intertemporal choice

In this study, we constructed a model in which a few different mechanisms could explain temporal discounting behavior, such as distorted encoding of the stimuli, attention prioritization to the delay information, or an inability to suppress the shorter sooner option. Fifteen different models were constructed from this overarching structure, each of which possessed a unique configurations of mechanisms. In addition to the core parameters *θ*, *τ* , and *σ*, our current study revealed that model performance was best achieved when the following three additional parameters were allowed to freely vary across subjects: *α*_*t*_, *β*_*I*_ , and *β*_*D*_.

In a previous application [15], we used a similar model to account for behavioral data from another intertemporal choice task. In a similar analysis, a factorial model fitting exercise suggested that model performance was best achieved when *ω*, *β*_*I*_ , and *β*_*D*_ were freely estimated – similar to Model 13 presented here. The consensus of the lateral inhibition terms *β*_*I*_ and *β*_*D*_ across the two studies suggest that these mechanisms are strong contributors in explaining valuation accumulation dynamics within intertemporal choice. By contrast, we can speculate about the disagreement between *ω* and *α*_*t*_ on two grounds.

First, in contrast to the design presented in [15], our current experimental design showed the time delay and amount of reward for the delayed option sequentially. Here, we adjusted the model to account for the sequential structure by first allowing subjective valuation of the alternatives to accumulate for the first second on the basis of only delay information. Once the reward information was presented, the model performed identically to that presented in [15]. Additionally, when parameters are not freely estimated, a choice must be made as to what value these fixed parameters should take. In [15], *ω* was set to 0.5, where attention is equally divided between the two dimensions. Here, we set *ω* to 0.9 in light of the potential asymmetry that might ensue, due to the delay information being presented first, and in isolation. Although we also tested *ω* = 0.5, with negligible difference in model performance, it is possible that these fixed parameter choices could at least partially explain the differences between the results. Second, it is also possible that choosing between a delayed option and a fixed immediate option in our study, and choosing between two delayed options (i.e., a shorter sooner and a larger later option) in [15], can partially induce the use of different mechanisms in the choice process.

### Frontoparietal functional connectivity

The present study identified that phase-based frontoparietal functional connectivity between putative dmFC and pPC was associated with the degree of competition occurring in the value accumulation process. This finding is in line with previous studies where EEG gamma band activity reflected the gradual accumulation of evidence in value-based decision making tasks [44]. [45] adopted simultaneous EEG–fMRI approach and identified the posterior-medial frontal cortex (PMFC) as the spatial correlates of such accumulation mechanisms. In addition, they found that the PMFC exhibited task-dependent coupling with the vmPFC and the striatum. Our study further endorsed the proposition of an evidence accumulation process during value-based decision making, and extended it with a choice task modulated by higher level of self-control rather than a preference-based hedonic decision making task in [45]. As such, our study postulated that the evidence accumulation mechanism may subserve a variety of decision making tasks. Also, the phase-based functional connectivity that we identified from EEG was between putative dmFC and putative pPC, extending from previous focus on within-frontal connectivity dynamics. Although it may be problematic to directly compare the spatial correlates of accumulation dynamics across different studies due to the different task structure and different modality of neural measures, some consensus between the two results is assuring. Future studies may shed light on the connections between these different studies.

Previous investigations of connectivity dynamics in intertemporal choice mostly relied on fMRI. For example, [12,46] investigated the connectivity between vmPFC and dlPFC, as evidence for the mechanism of self-control that the dlPFC has on the putative valuation region vmPFC. [13] identified functional connectivity between the vmPFC and three regions of dlPFC, bilateral pPC and dmFC, as a partial support for the value accumulation mechanism of the three regions. Our study was unable to localize the vmPFC or the ventral striatum due to the low spatial resolution of EEG measurements for sites distal from the scalp. Instead, our results add to our knowledge about how frontoparietal connectivity relates to choice competition in the accumulation process. Across these results, is not yet clear whether the connectivity between the dmFC and the pPC reflects a genuine connection between the two anatomical regions, or a common connection with vmPFC. Interestingly, we did not find such phase-based connectivity between dmFC and dlPFC.

### Links between brain and behavior

In this study, we constructed value accumulation dynamics from a computational model and related them to connectivity dynamics in our EEG data. Such a link between the two modalities is considered purely theoretical [47], and is relatively weak from the perspective of statistical power. A more integrated approach would be to use the neural dynamics to directly drive the computational model [48–51], or to use the covariation betwen their trial-to-trial fluctuations to modulate and constrain the compuational model [52,53]. Although an ideal neural measure would have trial-level information to substantiate mechanistic claims [15, 54–56], functional connectivity measured by ISPC requires several trials to establish a reliable measure, making a trial-level analytic approach infeasible for such precise linking in our study. Another possibility would be to establish a link to the computational model at the moment-by-moment level, which would further exploit the temporal resolution of EEG data provided by EEG measures. For example, a time-frequency analysis on the reconstructed accumulation trajectories might reveal similar information with respect to time and frequency, compared to a time-frequency result in ISPC.

Although providing high temporal resolution, EEG suffers from limited spatial resolution. The dmFC and pPC electrodes we identified provide tentative connections of the anatomical regions. However, it is difficult or impossible to establish connectivity results for other important deep structures or subcortical regions, such as vmPFC and ventral striatum. Therefore, the previous finding of connectivity between vmPFC and dlPFC cannot be replicated from purely EEG analysis. Due to the limitations of both approaches, a combination of EEG and fMRI, or even more modalities could provide a workaround on the measurement limitations and offer complementary correlates of neural activity. The idea of fusing EEG and fMRI is not novel, and there have been attempts using simultaneous fMRI and EEG on various decision making tasks including intertemporal choice (e.g. [57–59]. We argue that including cognitive representations for the behavioral data will also be an important connective medium, which has been discussed in detail elsewhere [56,60–63].

### Beyond hyperbolic discounting

Our model assumed that delay and reward amount are evaluated separately and are combined through direct comparison before being integrated into subjective values. This contrasts starkly with the standard mechanistic view of intertemporal choice (e.g. [64,65]). The standard model posits that subjective value is first calculated for both options using hyperbolic discounting. Decision making follows through comparison of subjective values. Daniel Read and colleagues have challenged this view [66] by pointing out that discounting across a time delay is not consistent with discounting across subintervals of that delay (i.e. discounting between 0 and *t*′ and between *t*′ and *t* can be greater than simply between 0 and *t*, where 0 < *t*′ < *t*, a phenomenon known as subadditivity). Scholten and Read [19] have proposed a new model of discounting that accounts for these effects and that simultaneously accommodates hyperbolic discounting. Their model is mechanistically similar to the model presented here, and we have also shown that our model provides excellent fits to hyperbolic discounting patterns [15]. A further significant advantage of our feature-based process model is that it provides a means to incorporate order dependence in intertemporal preference. In this experiment we showed delay information ahead of reward amount and our findings suggest that participants are influenced by this asynchrony so that decision making begins from biased starting points once reward information is presented, corresponding to well documented order effects in intertemporal choice (e.g., [67]). A next step for our modeling efforts will be to extend the generality of phenomena that feature-based process models accommodate.

## Methods

### Subjects

A total of 25 healthy adults participated in this study (12 females, ages 19-35 years, median 22 years). All subjects gave written informed consent before completing the experiment and all procedures were approved by Stanford University’s Institutional Review Board. Two subjects were excluded due to data collection problems resulting in excessive artifacts in the EEG data. Data from the remaining 23 subjects were analyzed (12 females, ages 19-35 years, median 22 years).

### Experimental design

Subjects completed a pretest session and an EEG session of the intertemporal choice task, with EEG acquired only in the second session. A similar experimental design has been used in our previous fMRI studies [13–15]. The pretest session used a staircase procedure to measure each individual’s discount rate *k*, assuming a hyperbolic discounting function,

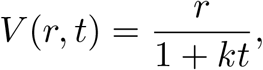

where *V* is the subjective value of a reward (in $), *r* is the monetary amount offered (in $), and *t* is the delay (in days; *t* = 0 for payment available “Today”). After completing the behavioral pretest task, we fit a softmax decision function to subjects’ choices. We assumed that the likelihood of choosing the delayed reward (*P*_*D*_) was given by a softmax rule,

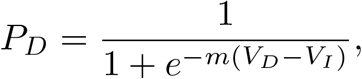

where *m* accounts for individual sensitivity to changes in discounted value, *V*_*I*_ is the value of the smaller sooner option (in $, *t* = 0 for all choices in this task), and *V*_*D*_ is the value of the larger later option. Using a maximum likelihood procedure, decision parameters (*k* and *m*) were estimated from the data obtained during the staircase task and used to generate stimuli for the EEG session. In this way, we were able to systematically manipulate the expected probability of choosing the delayed reward (*P*_*D*_) as the within-subject independent variable for the EEG session.

The staircase procedure used during the first task required subjects to select between a delayed reward (of *r* dollars available at delay *t*) and a fixed immediate reward of $10 (*V*_*I*_). For any choice, indifference between the immediate and delayed options implies a discount rate of *k* = (*r* − *V*_*I*_)(*V*_*I*_*t*). We refer to this implied equivalence point as *k*_*eq*_. Our procedure amounted to varying *k*_*eq*_ until indifference was reached. Specifically, we began with *k*_*eq*_ = 0.02, and if the subject chose the delayed reward, *k*_*eq*_ decreased by a step size of 0.01 for the next trial. Otherwise, *k*_*eq*_ increased by the same amount. Every time the subject chose both a delayed and an immediate offer within five consecutive trials, the step size was reduced by 5%. Subjects completed 60 trials of this procedure, and we placed no limits on the response time in this pretest session.

In the EEG session, we selected a 1000 ms separation between the presentation of delay and amount information to maintain a minimal epoch length that allowed subjects to make accurate intertemporal decisions. We wished to minimize the epoch length to ensure subject’s continual engagement and accurate time-locked signals in the EEG data. The 1000 ms separation was sufficient to allow us to distinguish between neural responses to delay and amount presentations with EEG. In addition, the sequential design allowed us to keep subjects fixated on the center of the screen, minimizing eye movement artifacts. After removing trials with excessive eye movements, the median number of trials across subjects was 179. Except for one subject left with only 115 trials after trial rejection (indexed as subject 15 in Fig 3c), trial numbers for other subjects were in the range of 168-180.

### EEG recordings and preprocessing

EEG data was collected using a 128 channel Geodesic Sensor Net (Electrical Geodesics, Inc., Eugene OR, USA), with a 500 Hz sampling rate, using the vertex as reference. During pre-processing, we re-referenced to the average reference, epoched trials from −1500 to +6500 ms around the onset of delay presentation, baseline corrected trials using the average from 0 to 5000 ms, and band-pass filtered the data at 0.5 - 200 Hz. Trials were visually inspected and rejected if excessive artifacts were present. Normally occurring artifacts were rejected using an independent component analysis algorithm from the EEGLab toolbox [68]. Epochs were then transformed to current source density (CSD) using the CSD toolbox [69]. CSD transformations of EEG data reduce the influence of volume conduction across the scalp and constrain the effect of cortical sources to within ≤ 5*cm*^2^ from the given electrode, facilitating the analysis of oscillatory neural activity and localization of activity from recordings at the scalp [70,71].

### Intersite phase clustering

We analyzed functional connectivity in the EEG data by performing statistical tests of intersite phase clustering (ISPC). ISPC quantifies the time-dependent connectivity strengths between putative brain regions by measuring the consistency of the difference of phase angles between a pair of electrodes [42]. To obtain ISPC from our EEG data, we convolved CSD-EEG epochs with a set of Morlet wavelets, which are defined as 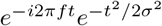 where *t* is time, *f* is frequency, and *σ* defines the width of each frequency band in terms of cycles. We constructed wavelets for 30 frequencies, from 2 – 30Hz in logarithmic steps, and varied the width for each of these wavelets, from 6 – 10 cycles, also in logarithmic steps. We computed ISPC between pairwise electrodes by extracting the phase angle (*ϕ*) from the convolved signal, and then calculated

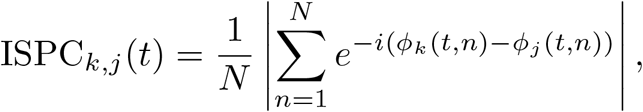

where *N* is the number of trials and *ϕ*_*k*_(*t, n*) is the instantaneous phase angle for electrode *k* in trial *n* at time *t*. This equation provides a measure for the consistency of the difference of phase angles between a pair of electrodes *k* and *j*. A perfectly consistent difference would result in a value of 1 in the equation above and imply strong functional connectivity, independent of amplitude co-variation [72] Uniformly distributed differences would produce a value of 0 and suggest lack of functional connectivity between two brain regions.

The above ISPC measure can be standardized as pISPC, by calculating the percent change in ISPC with reference to the grand average ISPC over the time window of interest within every frequency band, such that

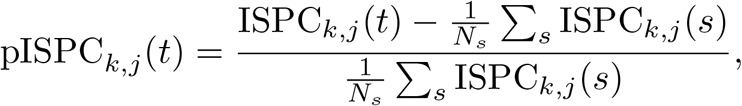

where the average ISPC is calculated within the time range of interest, denoted by *s*, and *N*_*s*_ denotes the number of time points in the time range. Therefore, pISPC provides us with a relative measure of increases in functional connectivity that can be compared between experimental conditions, for any time window and frequency band. Our analyses spanned the frequency range of 2 - 30 Hz in logarithmic steps and the time window of interest is from 0 - 5 seconds from the onset of delay presentation. The time window from 0 to 5 seconds is used as the reference to calculate pISPC. Compared with the convolved time windows from −1.5 to +6 seconds, the ISPC analyses have ignored the edges of the convolved time windows and so should not reflect edge artifacts.

### Model parameter estimation

We fit our behavioral model to individual data in a hierarchical Bayesian framework. For simplicity and based on our previous analyses, model parameters at the subject level were assumed to be distributed according to a normal distribution with group means and standard deviations as the shape parameters. Specifically, we assumed that for each subject *j*,

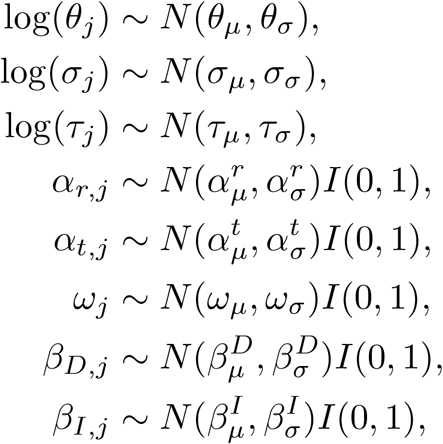

where *I*(*a, b*) denotes an indicator function on the interval (*a, b*). For the group mean parameters, we specified the following informative priors after ensuring that the prior predictive distributions created reasonable ranges for subject-level parameter values:

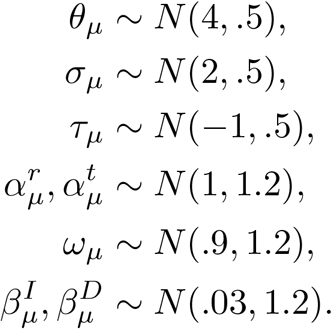

We adopted similarly informative priors for the group-level standard deviation parameters, such that

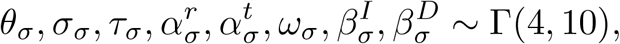

where Γ(*a, b*) indicates the Gamma distribution with shape parameter *a*, and rate parameter *b*.

When fitting each model configuration to behavioral data, due to the analytically intractable stochastic process, we approximated the likelihood function using likelihood-free Bayesian estimation techniques [73–75]; namely, we used the probability density approximation (PDA; [76]) method. The PDA method was nested within a Gibbs ABC algorithm [77] to allow for likelihood-free approximations of the subject-level parameters, but direct posterior sampling of group-level parameters. To facilitate efficient sampling of the joint posterior distribution, we used differential evolution with Markov chain Monte Carlo (DE-MCMC; [78,79]). We used 24 concurrent chains for 1000 iterations, following a burn-in period of 1000 iterations, resulting in 24000 samples of the joint posterior distribution. A migration step [79,80] was used with probability 0.1 for the first 250 iterations, after which time the migration step was terminated. We also used a purification step every 10 iterations to ensure that the chains were not stuck in spuriously high regions of the approximate posterior distribution [81]. Visual inspection was performed on the chains to ensure that chains converged well.

### Parameter recovery

We carried out a recovery study, showing in S2 Fig that our modeling fitting procedure was able to provide a good recovery for all parameters from Model 11 (the best performing model), over a wide range of values that correspond to individual MAP estimates from the experimental data. In particular, we simulated the synthetic data for each subject using the MAP estimates from Model 11. The MAP estimates were practically calculated as the average values of the last 100 iterations from the Markov chains. Given the synthetic data of each subject, we followed the same procedures to recover the model parameter, as were performed for actual experimental data.

## Supporting information

**S1 Fig. Distribution of Maximum a posteriori (MAP) parameter estimates across subjects from Model 11 (the best performing model).** Each panel corresponds to one of the six free parameters (*θ*, *σ*, *τ* , *α*_*t*_, *β*_*I*_ , and *β*_*D*_) in Model *θ*, *σ*, and *τ* are constrained to be positive values whereas *α*_*t*_, *β*_*I*_ , and *β*_*D*_ are constrained to be between 0 and 1, under the estimation procedure described in Methods section. See S1 Table for detailed values of individual parameter estimates.

**S2 Fig. Parameters recovery.** Each panel in this figure shows the recovered individual parameter values from synthetic data against the estimated parameter values from actual data. The six parameters are the freely estimated parameters in Model 11 (the best performing model). Each data point in each panel indicates one single subject, where *x*-axis corresponds to the MAP parameter estimates from actual data and *y*-axis represents the MAP parameter estimates from the synthetic data. We calculated the Pearson correlation coefficient for each parameter between the estimated parameter values and recovered parameter values and showed in each panel. For visual clarity, we excluded one outlier parameter value from one subject (subject 14).

**S3 Fig. Posterior prediction of choice and response time distributions from Model 11 (the best performing model).** Five panels correspond to the five *P*_*D*_ experimental conditions. For each panel, the histograms denote the observed data, where response times associated with immediate choices were plotted on the negative x-axis and response times associated with delayed choices were plotted on the positive x-axis. The overlaid black lines denote the simulated data, with simulated response times separated according to the simulated choices similarly. The comparison of experimental data and simulated data endorses the successful model fit of Model 11.

**S4 Fig. Posterior prediction of choice and response time distributions and summary statistics from Model 14 (the second best performing model).** This figure presents similar model performance summaries as we did for Model 11. The summary statistics (left) corresponds to Fig 5a for Model 11, and the choice and response time distribution (right) corresponds to S3 Fig.

**S1 Table. Maximum a posteriori (MAP) individual parameter estimates from Model 11 (the best performing model).** The listed six parameters were freely estimated in Model 11, with other parameters as fixed values: *α*_*r*_ = 1, *ω* = 0.9. Note that parameters *θ*, *σ*, and *τ* were estimated in the logarithmic space, and the MAP values presented in the table have been transformed back to their respective original space.

**S2 Table. Maximum a posteriori (MAP) individual parameter estimates from Model 14 (the second best performing model).** The listed six parameters were freely estimated in Model 11, with another parameter *ω* fixed as 0.9. Note that parameters *θ*, *σ*, and *τ* were estimated in the logarithmic space, and the MAP values presented in the table have been transformed back to their respective original space.

## Acknowledgments

The authors would like to thank Jerome Busemeyer for comments that improved an earlier version of this manuscript, Mike X Cohen for guidance on the EEG data analyses, and Tony Norcia for guidance on the design of the EEG experiment.

